# Functional Characterization of Calmodulin-like Proteins, CML13 and CML14, as Novel Light Chains of Arabidopsis Class VIII Myosins

**DOI:** 10.1101/2023.05.12.540561

**Authors:** Kyle Symonds, Howard J. Teresinski, Bryan Hau, Einat Sadot, Vikas Dwivedi, Eduard Belausov, Sefi Bar-Sinai, Motoki Tominaga, Takeshi Haraguchi, Kohji Ito, Wayne A. Snedden

## Abstract

Myosins are important motor proteins that associate with the actin cytoskeleton. Structurally, myosins function as heteromeric complexes where smaller light chains, such as calmodulin (CaM), bind to isoleucine-glutamine (IQ) domains in the neck regions to facilitate mechano-enzymatic activity. We recently identified Arabidopsis CaM-like (CML) proteins, CML13 and CML14 as interactors of proteins containing multiple IQ domains, including a member of the myosin VIII class. Here, using *in vivo* and *in vitro* assays we demonstrate that CaM, CML13, and CML14 bind the neck region of all four Arabidopsis myosin VIII isoforms. Among ten CML isoforms tested for *in planta* binding to myosins VIIIs, CaM, CML13, and CML14 gave the strongest signals using *in planta* split-luciferase protein-interaction assays. *In vitro,* recombinant CaM, CML13, and CML14 showed specific, high-affinity, calcium-independent binding to the IQ domains of myosin VIIIs. Subcellular localization analysis indicated that CaM, CML13, and CML14 co-localized to plasma membrane-bound puncta when co-expressed with RFP-myosin fusion proteins containing IQ- and tail-domains of myosin VIIIs. In addition, *in vitro* actin-motility assays using recombinant myosin holoenzymes demonstrated that CaM, CML13, and CML14 function as light chains for myosin VIIIs. Collectively, our data indicate that Arabidopsis CML13 and CML14 are novel myosin VIII light chains.

**Highlight:** Myosins are key proteins in the plant cytoskeleton, but the identity of their light chain components is unknown. Here, we show that calmodulin-like proteins function as novel myosin light chains.

## INTRODUCTION

The cytoskeleton is a dynamic and complex system that functions in the organization of the intracellular environment. A ubiquitous component of the cytoskeleton is the acto-myosin network that contains myosin motor proteins bound to actin polymers. The myosin superfamily is broadly organized into two groups: conventional and unconventional. Conventional myosins form filaments and mainly act in animal muscle contraction and cell motility (Altman, 2013). In contrast, unconventional myosins do not form filaments and are involved in various processes such as organelle trafficking, cell and organism growth, and nuclear rearrangements (Hartman *et al*., 2011; Altman, 2013; Nebenführ and Dixit, 2018). These unconventional myosins possess four distinct domains: the motor, neck, coiled-coil, and tail domains.

Although there are 79 phylogenetic classes of myosins (Kollmar et al., 2017), only of these, VIII and XI, are present in plants (Nebenführ and Dixit, 2018). The Arabidopsis genome encodes four class VIII and thirteen class XI myosins, and studies indicate a large degree of functional overlap among these (Avisar et al., 2009; Peremyslov *et al*., 2010; Haraguchi *et al*., 2018). Myosin XIs are involved in a range of processes including endo and exocytosis, cytoplasmic streaming, nuclear shape and positioning, gravitropism, and biotic stress responses (Peremyslov *et al*., 2010; Tominaga and Nakano, 2012; Buchnik et al., 2015; Nebenführ and Dixit, 2018). In contrast, the roles of myosin VIIIs are not as well understood and a quadruple knockout of all four myosin VIII genes in Arabidopsis shows only subtle phenotypes (Talts et al., 2016; Liu *et al*., 2023). Nevertheless, studies on class VIII myosins have implicated them in plasmodesmatal function and endocytosis (Baluška et al., 2001; Golomb et al., 2008; Sattarzadeh et al., 2008). Their enzymatic profile suggests tension sensor and/or tension generator activity rather than fast movement (Haraguchi et al., 2014; Henn and Sadot, 2014; Rula et al., 2018). In addition, recent reports show the involvement of myosin VIIIs in the movement of viral particles of tobacco mosaic, rice stripe, and rice grassy stunt viruses and the VirE2 protein from *Agrobacterium tumefaciens* (Pitzalis and Heinlein, 2017; Liu *et al*., 2023). In the moss *Physcomitrella patens*, myosin VIIIs function during cytokinesis (Wu and Bezanilla, 2014).

In a generalized model of myosin architecture, the catalytic motor (head) domain of myosins interacts with actin filaments and is responsible for ATP hydrolysis, the neck domain binds one or more myosin light chains (MLCs) via IQ (isoleucine/glutamine) motifs, the coiled-coil domain allows for homodimerization, and the globular tail domain binds to various cargo (Peremyslov *et al*., 2013; Kurth *et al*., 2017; Duan and Tominaga, 2018). Very little is known about the neck domains of plant myosins, but they serve as the lever arm during mobility and are thus essential for proper myosin function. Myosin necks possess one or more IQ domains, defined by the general consensus motif IQXXXRGXXXR, that are arranged in close proximity and act as the binding sites for MLCs that provide the rigidity essential for myosin movement (Heissler and Sellers, 2014). Plant myosin VIIIs are predicted to possess 3-4 IQ domains whereas myosin XIs have 5-6 (Nebenführ and Dixit R, 2018). MLCs are thus critical myosin partners and components of the cytoskeleton network. Historically, MLCs have been referred to by various names however, upon their discovery, the MLC occupying the first IQ motif of muscle myosin II was called the “essential” light chain (ELC). The term “essential” was chosen as the ELC could only be removed under harsh conditions and dissociation inactivated the holoenzyme (Heissler and Sellers, 2014). Similarly, the MLC bound to the second IQ motif was termed the “regulatory” light chain (RLC), as this MLC was involved in regulating myosin II’s motor activity (Lee *et al*., 2022). MLCs described to date are either the evolutionarily-conserved, Ca^2+^-binding protein calmodulin (CaM) or are members of the EF-hand-containing superfamily (Heissler and Sellers, 2014).

It is thought that Ca^2+^ modulates acto-myosin dynamics in plants by decreasing myosin XI motility in response to an increased Ca^2+^ concentration (Yokota et al., 1999), but the model is based primarily on studies of mammalian class V myosins (Tominaga and Nakano, 2012). MLCs such as CaM regulate class V myosins by binding to the IQ domains of the neck region in a Ca^2+^-independent manner and Ca^2+^-induced dissociation of MLCs from the neck domain *in vivo* is thought to be the main mode of regulation (Manceva *et al*., 2007; Tominaga and Nakano, 2012). In this respect, IQ domains are distinct from most other (non-IQ) CaM-binding domains where CaM binds in the Ca^2+^-bound (holo-CaM) form (Clapham, 2007). CaM is well-established as an MLC in animals, and myosins from both lily pollen tubes and a green algae (*Chara corallina*) co-purified with CaM and other, unidentified, Ca^2+^-binding, non-CaM proteins that may be alternative MLCs (Kakei *et al*., 2012). This suggests that CaM acts as a light chain for some plant myosins *in vivo*. However, some recombinant Arabidopsis myosin XIs co-expressed solely with CaM in insect cells do not exhibit smooth motility *in vitro* (Haraguchi *et al*., 2018), suggesting that, in addition to CaM, other plant-specific MLCs may be needed for mechanoenzyme function. By comparison, various non-CaM MLCs have been identified and characterized in mammalian and fungal species, where they often perform vital roles (Heissler and Sellers, 2014).

Despite their potential importance in myosin function, very little is known about plant MLCs and whether any of the CaM-related proteins found in plants may function as MLCs. In addition to the evolutionarily-conserved CaM, plants also possess a large family of unique CaM-like (CML) proteins, with 50 isoforms in Arabidopsis that range from about 20-80% sequence identity with CaM (McCormack and Braam, 2003; Zhu *et al*., 2015). The only functional domains in CMLs are Ca^2+^-binding EF-hands, and thus they are predicted to function like CaM as regulatory proteins via interaction with downstream targets. While the functions of most CMLs remain unclear, several have been shown to participate in abiotic and biotic stress responses as well as during various stages of development (DeFalco *et al*., 2009; La Verde *et al*., 2018). However, unlike CaM, where many targets have been characterized, most CML targets remain unknown (La Verde *et al*., 2018).

Several previous studies speculated that CaM and some CMLs may function as MLCs in plants. For example, CaM was proposed to be the ELC of myosin ATM1 (also known as myosin VIII-1) as it bound to IQ1 and increased the actin sliding velocity *in vitro* (Haraguchi *et al*., 2014). However, when additional IQs in the neck domain of ATM1 were included, the motility experiments were not successful in the presence of CaM alone, suggesting that additional MLCs are likely needed for the function of ATM1 and possibly other myosins (Haraguchi *et al*., 2014, 2018). Similarly, a screen for cytoskeletal modulators using fibroblast cells isolated CML24 (aka TCH2) as a putative interactor of the neck domain of ATM1 *in vitro* (Abu-Abied *et al*., 2006). Recently, proteome analysis of complexes associated with Arabidopsis SnRK1-α1 and -γ1 identified myosin XI isoforms as well as CML13 and CML14 (Van Leene *et al*., 2022). These studies provide circumstantial evidence that some CMLs may function as MLCs in plants. We recently reported that CML13 and CML14 interact with proteins that possess tandem IQ domains, including members of the IQ67-domain family (IQDs), CaM-binding transcriptional activators (CAMTAs), and class VIII myosins (Teresinski *et al*., 2023). Here, we expand on these previous findings to investigate whether CML13 and CML14 are functional MLCs. We focused on the four members of the Arabidopsis myosin VIII family and used a combination of *in vitro* and *in vivo* approaches to assess the interaction of these myosins with CaM and various CMLs. Our data strongly suggest that conserved CaM, as well as CML13 and CML14 function as novel MLCs in Arabidopsis.

## MATERIALS AND METHODS

### Plant material and growth conditions

For split-luciferase protein interaction assays, *Nicotiana benthamiana* seeds were sown in Sunshine mix #1 (Sun Gro Horticulture Canada Ltd.) and transferred to a growth chamber under short-day conditions (Conviron MTR30; 12h photoperiod, 22°C, ∼150 *μmol* m^-2^ s^-1^). After 14 days, individual seedlings were transplanted to independent 10-cm pots and returned to the growth chamber. Pots were sub-irrigated as needed, with N-P-K fertilizer (20-20-20, 1 g/L) applied every week. For fluorescence microscopy assays, *N. benthmaniana* growth conditions were as previously described (Belausov et al., 2023).

### Plasmid constructs and recombinant protein expression

For PCR and cloning, oligonucleotide primers used are listed in Supplementary Table S1. See Supplementary Table S2 for a description of plasmid constructs used in this study and the corresponding regions of proteins encoded by the respective cDNAs. The cDNAs encoding a representative group of CMLs (CML6, 8, 15, 19, 24, 35, 38, and 42), as well as CML13, CML14, and CaM81 were cloned as full-length, open-reading frames into the pCambia1300-C-Luciferase (CLuc) vector downstream of the firefly luciferase enzyme as described previously (Chen et al., 2008). CaM81 (hereafter, referred to as CaM) is a conserved isoform of CaM from petunia (Genbank accession M80836) that is identical at the protein level to Arabidopsis CaM7. cDNAs encoding the full region or truncations of the *Arabidopsis* myosin class VIII neck domains were cloned upstream of the N-terminus of firefly luciferase in the pCambia1300-N-Luciferase (NLuc) binary vector. For fluorescent protein fusions we used the Golden Gate system (Engler et al., 2014). The cDNAs of CML13, CML14, CaM with flanking adapters having a BsaI site (5’ TGGTCTCA**AATG** and 3’ GG**TTCG**TGAGACCA) and lacking the stop codon were synthesized by Twist Bioscience. The cDNA of GFP containing a linker sequence at the 5’ end (GGATCAACGGGTTCT encoding 5AA-GSTGS), was synthesized by Twist Bioscience. The BsaI adapters for GFP marker were TGGTCTCA**TTCG** at 5’ end and **GCTT**TGAGACCA at 3’ end. The plasmids encoding for 35Sx2pro and 35S terminator were pich41295 and pich41276, respectively. All the level-0 plasmids were mixed to create a level-1 backbone binary plasmid (pICH47742) by cut and ligate reaction using BsaI restriction enzyme and T4 DNA ligase from New England BioLabs NEB. All plasmid constructs were confirmed by DNA sequencing. The plasmid encoding the RFP–ATM1, RFP– ATM2, RFP–VIIIA and RFP–VIIIB were previously described (Avisar et al., 2009). *Agrobacterium* infiltration was performed as described (Belausov et al., 2023).

For recombinant protein expression, the cDNA sequences of interest were subcloned into expression vectors (Supplementary Table S2). cDNAs encoding the neck regions of myosin ATM1 (At3g19960) and ATM2 (At5g54280) were cloned into pET28b, pET28b-GB1, and pGEX-4T-3 vectors (Novagen) for expression in *E. coli* strain BL21 (DE3) CPRIL (Novagen). Recombinant proteins corresponding to the full-neck regions of myosin VIIIs were solubilized using 8 M urea and purified using a Ni-NTA His60 column (Takara), or for GST-tandem-IQ fusions, using a glutathione sepharose (Sigma) column, as per manufacturer’s instructions. Myosin full-neck proteins were maintained in 8 M urea as they were unstable in aqueous buffers, whereas proteins comprised of tandem IQ domains were dialyzed overnight against tris-buffered saline (TBS; 50 mM Tris-HCl, 150 mM NaCl, pH 7.5). Recombinant CaM, CML13, and CML14 were expressed and purified as described (Teresinski *et al*., 2023).

For the myosin *in vitro* sliding assays, the ATM1 construct encodes the motor domain and native neck regions (four IQ motifs) and the ATM2 construct encodes a motor domain and native neck regions (three IQ motifs) of ATM2. Full-length cDNA of ATM1 (At3g19960) and ATM2 (At5g54280) were provided by the RIKEN BioResource Center and Dr. Motiko Tominaga, respectively. A baculovirus transfer vector for ATM1 and ATM2 were generated using PCR (Supplementary Table S1). PCR products were ligated into the NcoI–AgeI restriction sites of pFastBac MD (Ito et al., 2009). The resulting constructs, pFastBac ATM1 and ATM2 encode an N-terminal amino acids (MDYKDDDDKRS) containing the FLAG tag (DYKDDDDK), amino acid residues 1–935 and 1–968 of ATM1 4IQ and ATM2 3IQ, respectively, and C-terminal amino acids (GGGEQKLISEEDLHHHHHHHHSRMDEKTTGWRGGHVVEGLAGELEQLRARLEHHPQGQRE PSR) containing a flexible linker (GGG), a Myc-epitope sequence (EQKLISEEDL), His tag (HHHHHHHH) and SBP tag (MDEKTTGWRGGHVVEGLAGELEQLRARLEHHPQGQREP). ATM1 and ATM2 were expressed using a baculovirus expression system in *High Five* insect cells. They were purified using nickel-affinity and FLAG-affinity resins as previously described (Haraguchi et al. 2022).

### Split-luciferase assay

The split-luciferase (SL) assays were performed as described (Chen et al., 2008). Six-week-old *N. benthamiana* leaves were infiltrated by *A. tumefaciens* strain GV3101. Whole leaves were removed after 4 d incubation and leaves were sprayed with a 1 mM D-luciferin (GoldBio) and 0.01% Silwett L-77 (Lehle Seeds Inc) solution and incubated in the dark for 10 mins. Leaves were imaged on the ChemiDoc™ Touch Imaging System (Biorad Inc.). Alternatively, leaf discs were taken after 4 d, incubated with 100 μL water containing 1 mM luciferin in a 96-well plate for 10-15 min, and luminescence was captured with the SpectraMax Paradigm multimode detection platform (Molecular Devices Inc.). The expression of C-Luciferase fusions was verified by immunoblot (Santa-Cruz Biotech).

### Microscopy and image analysis

For live cell imaging, a Leica SP8 confocal microscope was used. Imaging was performed using HyD detectors, HC PL APO CS 63x /1.2 water immersion objective (Leica, Wetzlar, Germany), and an OPSL 488 laser for GFP excitation with 500–530 nm emission range, and an OPSL 552 laser for RFP with 565–640 nm emission light detection. Colocalization analysis was done using the coloc function of Imaris (Oxford Instruments). The amount of 10 cells were analysed from each treatment. Rate of colocalization was calculated by the function of Pearson’s coefficient analysis in Imaris. Graphs and statistics analysis were performed by GraphPad Prism 9.5.1 using One-way Anova.

### Fluorescent protein-protein interaction overlay assays

CaM/CML-binding overlay assays were performed as previously described (DeFalco *et al*., 2010) with minor changes. Briefly, pure recombinant myosin VIII fusion proteins were blotted onto nitrocellulose membranes according to the figure legend and blocked overnight at 4°C with TBST (TBS and 0.01% Tween-20 (v/v)) supplemented with 5% Casein (w/v). The blots were washed and then incubated with 200 nM of pure, recombinant CaM or CMLs that had been covalently tagged with 680RD-NHS Ester (Li-Cor Biosciences) as per the manufacturer’s instructions. Protein-protein overlay assays were conducted in the presence of 2 mM CaCl_2_ or 5 mM EGTA for 1 hour at room temperature. The blots were washed several times in excess buffer without probe and imaged on the Odyssey XF Imager (Li-Cor Biosciences) on the 700 nm channel with a 30-second acquisition time.

### Steady-state dansyl fluorescence spectroscopy

Recombinant CaM and CMLs were covalently labeled using dansyl fluoride as described (Alaimo *et al*., 2013). Samples of 600 nM D-CaM or 3 µM CMLs were incubated in TBS with 1 mM CaCl_2_ or 1 mM EGTA with or without a 10-molar excess of IQ motif peptide, as indicated in the figure legend, for 30 mins at room temperature. Fluorescence spectra were collected using an excitation of 360 nm and emission wavelengths of 400-600 nm for CaM (Alaimo *et al*., 2013), and 400-650 nm for CML13 and CML14 at 25°C with a SpectaMax Paradigm instrument. Peptides were synthesized commercially by GenScript with the following sequences; ATM1-IQ1, LHGILRVQSSFRGYQARCLLKEL and ATM2-IQ1, LQGIVGLQKHFRGHLSRAYFQNM.

### Actin Gliding Assays

Actin-sliding velocity was measured using an anti-Myc antibody-based version of the *in vitro* actin-gliding assay as described (Ito et al. 2007). The velocity of actin filaments was measured in 25 mM KCl, 4 mM MgCl_2_, 25 mM Hepes-KOH (pH 7.4), 1 mM EGTA, 3 mM ATP, 10 mM DTT and oxygen scavenger system (120 μg/ml glucose oxidase, 12.8 mM glucose, and 20 μg/ml catalase) at 25°C. CaM, CML13, and CML14 alone or in different combinations of two putative MLCs were added in the assay buffer. Average sliding velocities were determined by measuring the displacement of actin filaments. Ionic strength was calculated using CALCON based on Goldstein’s algorithm (Goldstein 1979).

## RESULTS

### Sequence comparisons of Arabidopsis CaM, CML13, and CML14 and the myosin VIII neck domains

Comparisons of the primary structure of the neck regions of Arabidopsis myosin VIII isoforms are presented in Figure 1A and Arabidopsis CML13, CML14, and CaM in Figure 1B. A phylogenetic tree (Fig. 1C) based on the 17 Arabidopsis myosins shows the clear subgrouping of class VIII members. A sequence comparison and phylogenetic tree of CML13/14 orthologs from several plant taxa are presented (Supplemental Fig. 2) and are also compared to MLCs from a range of plant and nonplant species (Supplemental Fig. 3). Arabidopsis paralogs CML13 and CML14 are 95% identical and differ by only 8 residues while sharing ∼50% identity to CaM. Where CaM possesses four EF-hands, only the first EF-hand of CML13 and CML14 is predicted to be functional according to InterPro (Paysan-Lafosse *et al*., 2023) as there are gaps in EF2 and changes in key consensus residues in EF3 and EF4 relative to CaM (Fig. 1B).

**Figure 1.**
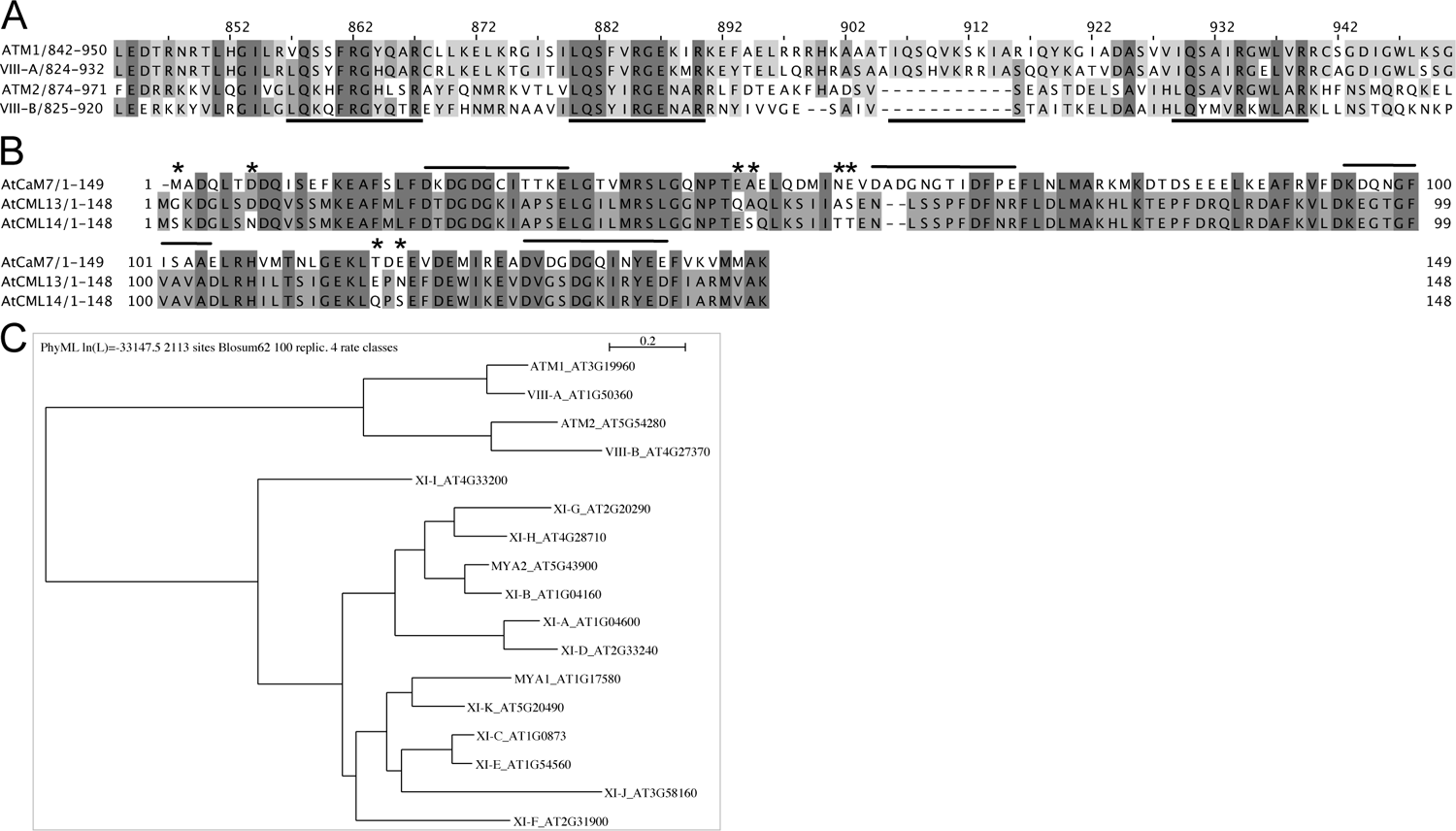
Protein sequence alignments of (A) the neck regions (residues ∼800-1000) for Arabidopsis class VIII myosins, and (B) Arabidopsis CaM, CML13, and CML14. Amino acid residues were shaded based on their percent identity, dark grey if identical, and progressively lighter grey until white as unconserved. In panel A, predicted IQ motif consensus sequences are underlined. The EF-hands of CaM in panel B are overlined and residues that differ between CML13 and CML14 are indicated with an asterisk. ClustalΩ was used for alignment (Sievers and Higgins, 2014; Gouy *et al*., 2021) and images were generated using Jalview Version 2.11.2.6. (C) Phylogenetic tree of the motor domains of Arabidopsis class VIII and XI myosins. The tree was constructed using Seaview version 5.0 (Sievers and Higgins, 2014; Gouy *et al*., 2021) from a complete alignment of proteins using neighbour-joining with bootstrapping analysis (1000 reiterations).

In the myosin VIII neck alignment, the core IQ consensus motifs are underlined and indicate that the neck regions of ATM1 and VIII-A, are each predicted to possess four IQ domains, whereas ATM2 and VIII-B possess three, having lost the IQ domain that corresponds to the third IQ-domain of ATM1 and VIII-A (Fig. 1A). IQ3 is quite degenerate in ATM1 and VIII-A and is missing several of the five core residues within the IQXXXRGXXXR canonical motif. Conversely, the most conserved IQ motif among myosin VIII isoforms, IQ2, deviates from the consensus myosin IQ sequence in that it possesses a negatively charged residue (Glu) at position 8 in place of hydrophobic residues which typically occupy this position (Houdusse *et al*., 2006). The most C-terminal IQ domain is relatively conserved among the isoforms and matches the typical CaM-binding IQ consensus, the exception being VIII-A which has a Glu in place of a large hydrophobic residue, (Fig. 1A). Given these sequence differences between CML13/14 and CaM, and among the IQ domains of myosin VIIIs, we sought to assess the interaction of these proteins and the potential of CaM, CML13, and CML14 to function as MLCs.

### CMLs bind the neck domain of Arabidopsis class VIII myosins *in planta*

To assess the association of Arabidopsis CaM and CMLs with the neck domains (i.e. the IQ-domain regions) of the myosin VIIIs, we used leaves of *N. benthamiana* and the split-luciferase (SL) protein-interaction assay (Fig. 2). A comparative schematic showing the domains of the myosin VIIIs is presented (Fig. 2A). Immunoblotting, using antisera against the C-terminus of firefly luciferase confirmed the expression of the CML-luciferase fusion constructs *in planta* (Supplemental Fig. S1). A representative full-leaf image of a SL assay testing interaction of ATM1 with CaM, CML13, CML14, and CML42 is presented in Supplementary Figure 4 and emphasizes the variability of the SL system as the level of bacterial infiltration, *Agrobacterial* infection efficiency, and transient protein expression are all aspects that cannot be controlled (Bashandy *et al*., 2015). Although common in the literature, empty vectors do not make suitable negative controls in the SL system. As such, we assigned CML42 a value of 1.0 as our base-level (negative) control, indicative of non-specific, background luciferase activity in the SL assay based on our previous study which indicated that CML42 does not associate with IQ domains (Teresinski *et al*., 2023). In pairwise tests, we observed the interaction of CaM with all four myosin VIII isoforms at a level about 3- to 4-fold (note log scale) above CML42 controls (Fig. 2B-E), consistent with reports that CaM is an MLC in plants (Ma and Yen, 1989; Yokota and Shimmen, 1994; Haraguchi *et al*., 2014, 2018). CML24 gave weaker signals, about 2-fold above background, with ATM1, VIII-A, and ATM2, but did not interact with VIII-B (Fig. 2B-E). By comparison, CML13 and CML14 showed interaction signals with ATM2 and VIII-B around 3- to 5-fold above controls, similar to those for CaM, but displayed markedly stronger signals with ATM1 and VIII-A, at levels about 35- and 200-fold, respectively, above CML42 controls (Fig. 2B-E). These observations corroborate a previous report of CML24 as a putative interactor of ATM1 (Abu-Abied *et al*., 2006) and demonstrate that CaM, CML13, and CML14 are able to bind to myosin VIIIs *in planta*.

**Figure 2.**
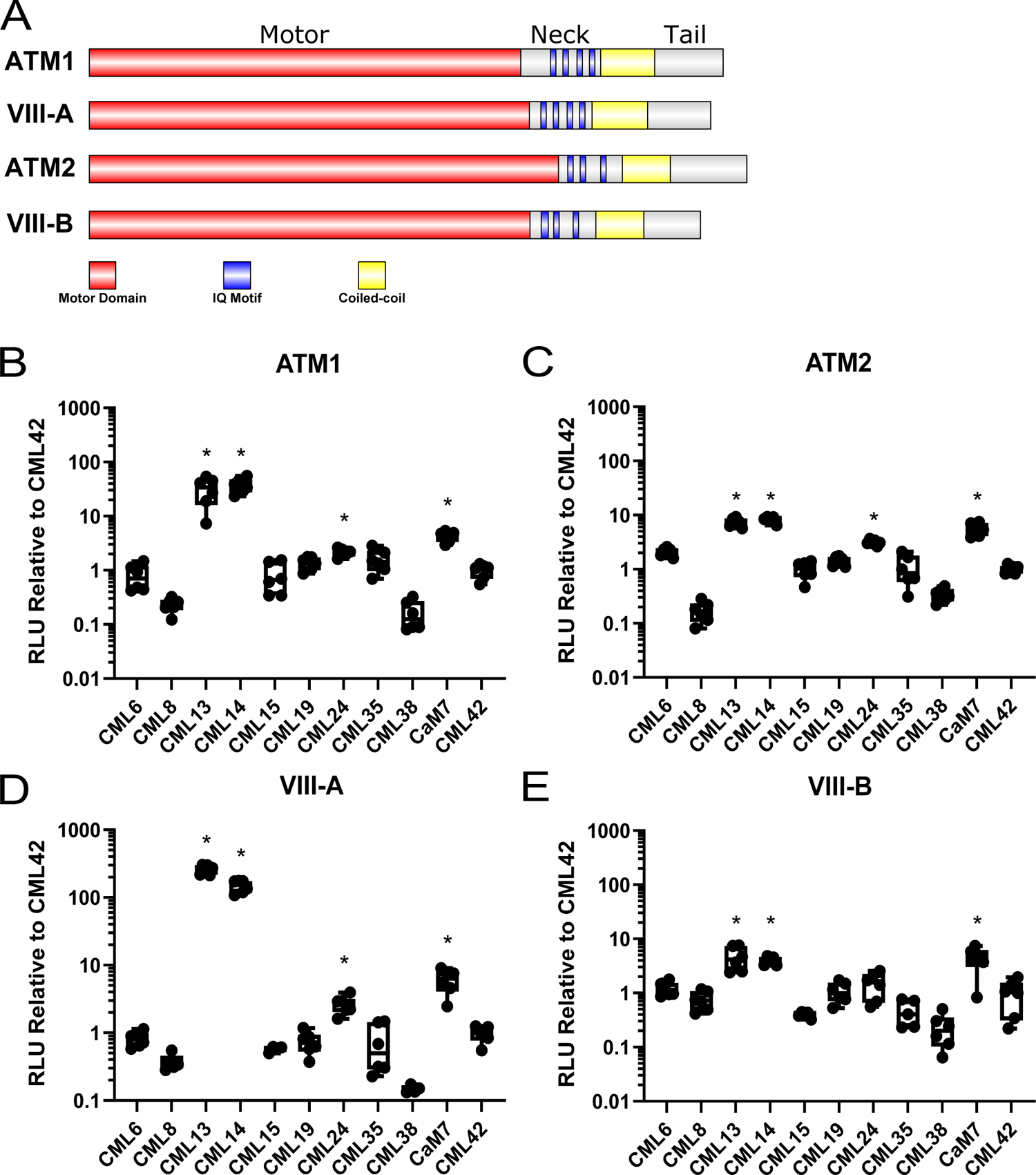
Split-luciferase protein interaction of Arabidopsis CMLs with the neck region of class VIII myosins *in planta*. *N. benthamiana* leaves were infiltrated with *Agrobacterium* harboring respective NLuc-prey (myosin VIII neck regions) and CLuc-bait (CaM, CMLs) vectors and tested for luciferase activity 4 days after inoculation, as described in Materials and Methods. (**A**) Schematic representation of myosin VIIIs showing the relative positions of myosin domains. *In planta* analysis of various CMLs with the neck regions of myosin (**B**) ATM1, (**C**) ATM2, (**D**) VIII-A, (**E**) VIII-B, respectively. Split-luciferase data are expressed as a fold-difference relative to the RLU signal (log_10_ scale) observed using the negative control bait CML42 which was set to an RLU of 1.0. Boxes contain each data point for 6 technical replicates, means are shown by a horizontal bar, the grey region is the 95% confidence interval, and whiskers extend to maximum and minimum data points. Asterisks indicate a significantly higher signal vs CLuc-CML42 as a negative control bait (One-way ANOVA against CML42 with Sidak’s test for multiple comparisons, p-value < 0.05). Data is representative of at least three independent experiments. RLU; relative light units.

We further explored the specificity of the myosin VIII/MLC interaction by testing the neck region of all four myosin VIIIs with representative isoforms from each of the nine CML subfamilies in Arabidopsis (CML6, 8, 13, 14, 15, 19, 24, 35, 38, with CML42 as a negative control) (Fig. 2B-E). Only CaM, CML13, CML14, and CML24 showed significant signals relative to CML42 controls. CML13 and CML14 typically gave the strongest signals in these protein-interaction tests. Taken together, these data indicate a clear specificity among the Arabidopsis CML family for interaction with myosin VIIIs, with CaM, CML13, CML14, and CML24 representing candidate MLCs.

### CML13 and CML14 co-localize to the plasma membrane with class VIII myosins

To test whether CML13, CML14, and CaM can co-localize with myosin VIIIs within cells, transient fluorescent confocal microscopy was performed. We used RFP fused to truncated myosin VIIIs that included the neck and tail region because the subcellular localization of these RFP-myosins has been previously reported (Golomb *et al*., 2008; Bar-Sinai *et al*., 2022). GFP-CaM or -CML constructs were transiently over-expressed in the presence or absence of RFP-myosin fusion constructs in *N. benthamiana* (Fig. 3). When expressed alone, we observed the localization of CML13 and CML14 to the cytoplasm whereas myosin VIIIs localized to discrete puncta at the plasma membrane, consistent with previous reports (Golomb *et al*., 2008; Teresinski *et al*., 2023). Interestingly, when GFP-CMLs and RFP-myosins were co-expressed, CML13 and CML14 distinctly altered their localization and were recruited to the punctate structures at the plasma membrane populated by the RFP-myosin fusion proteins (Fig. 3A, B) (Kumari *et al*., 2021). CaM, however, showed variable co-localization with myosin VIIIs, mainly co-localizing with ATM2 and to a lesser degree the other myosin VIII isoforms (Fig. 3C). Pearson’s coefficients of co-localization suggested that CML13 and CML14 interacted comparably with all myosin VIIIs except myosin VIII-B, which showed a lower coefficient relative to the other isoforms (Fig. 3D, and E).

**Figure 3.**
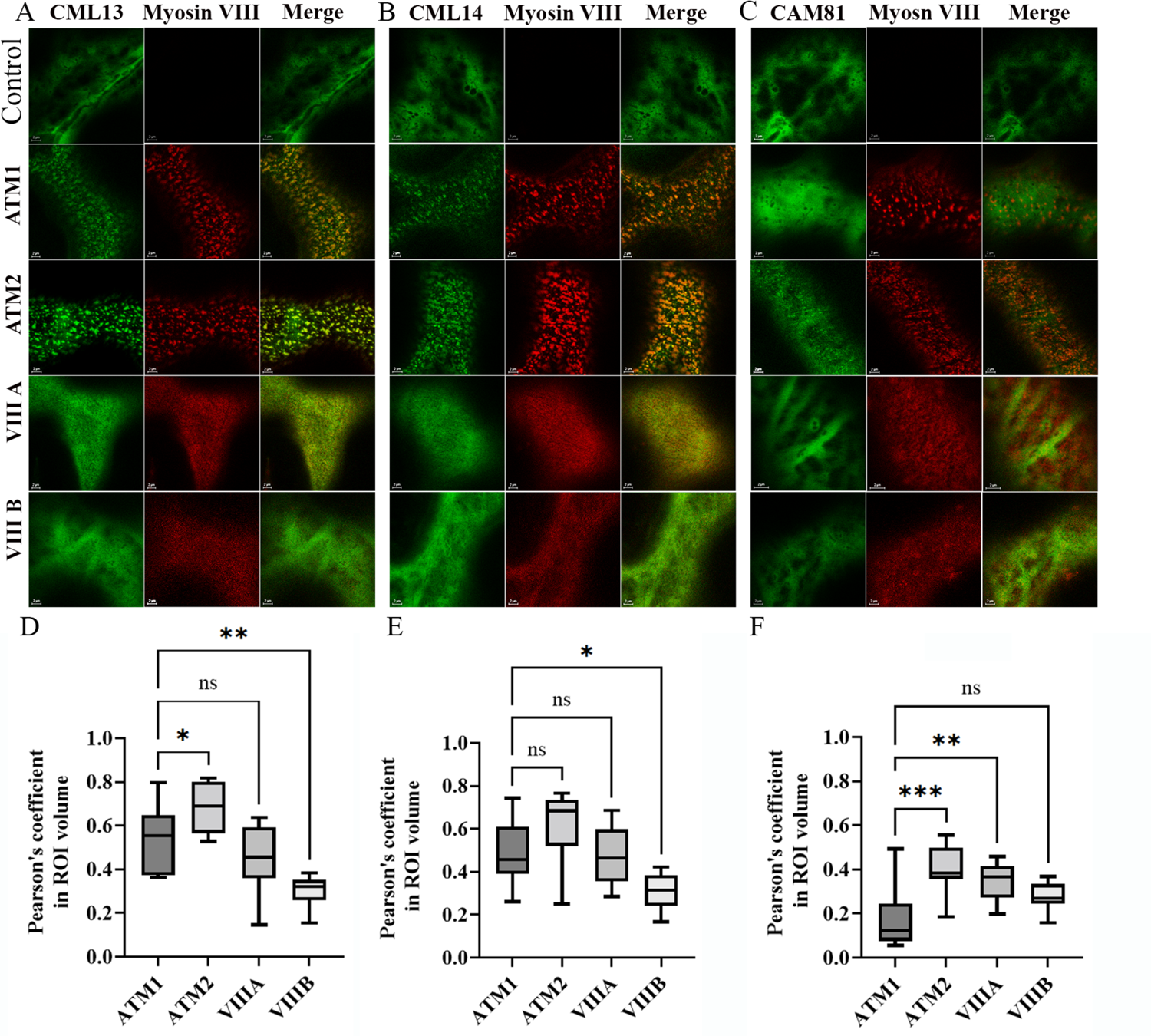
Colocalization analysis of myosin VIII IQ-tail fragments with CML13, CML14 and CAM81. RFP-ATM1^IQ-tail^, RFP-ATM2 ^IQ-tail^, RFP-VIII-A ^IQ-tail^, or RFP-VIIIB ^IQ-tail^ were transiently expressed in *N. benthamiana* leaves with (**A**) GFP-CML13 or (**B**) GFP-CML14 or (**C**) GFP-CAM81. Microscopy was performed 48h after Agro-infiltration using Leica SP8 confocal microscope. Pearson’s coefficient analysis for colocalization a was done (D) for GFP-CML13, (E) GFP-CML14 and (F) GFP-CAM81. Significance of differences was analysed by One-way Anova *p<0.05, **p<0.01, ***p<0.001

### Class VIII myosin IQ motifs have different specificities for CML13, CML14, and CaM

As CaM, CML13, and CML14 gave the strongest interaction signals in the SL system, we explored whether they exhibit specificity for distinct IQ domains within a given myosin. ATM1 and ATM2 were chosen as representatives among the two class VIII subgroups (Fig. 1C) and their neck domains were sequentially truncated and tested in SL assays for interaction with CaM, CML13, and CML14. Data were again expressed relative to CML42 as the negative control. Myosin IQ-motif pairs, which are thought to facilitate cooperative MLC interactions in other myosins (Langelaan *et al*., 2016; Pazicky *et al*., 2020), as well as single IQ motifs were tested for interaction with the putative MLCs. For ATM1, the tandem IQ pairs, ATM1-IQ1+2 and ATM1-IQ3+4, and the singles IQ domains, ATM1-IQ3 and ATM1-IQ4, bound to CML13, CML14, and CaM (Fig. 4A). ATM1-IQ1 gave the strongest signal with CaM in the SL assays, suggesting a possible preference for CaM over CML13 and CML14. In contrast to ATM1-IQ1, ATM1-IQ2 when tested alone showed statistically significant interaction exclusively with CML13 (Fig. 4A).

**Figure 4.**
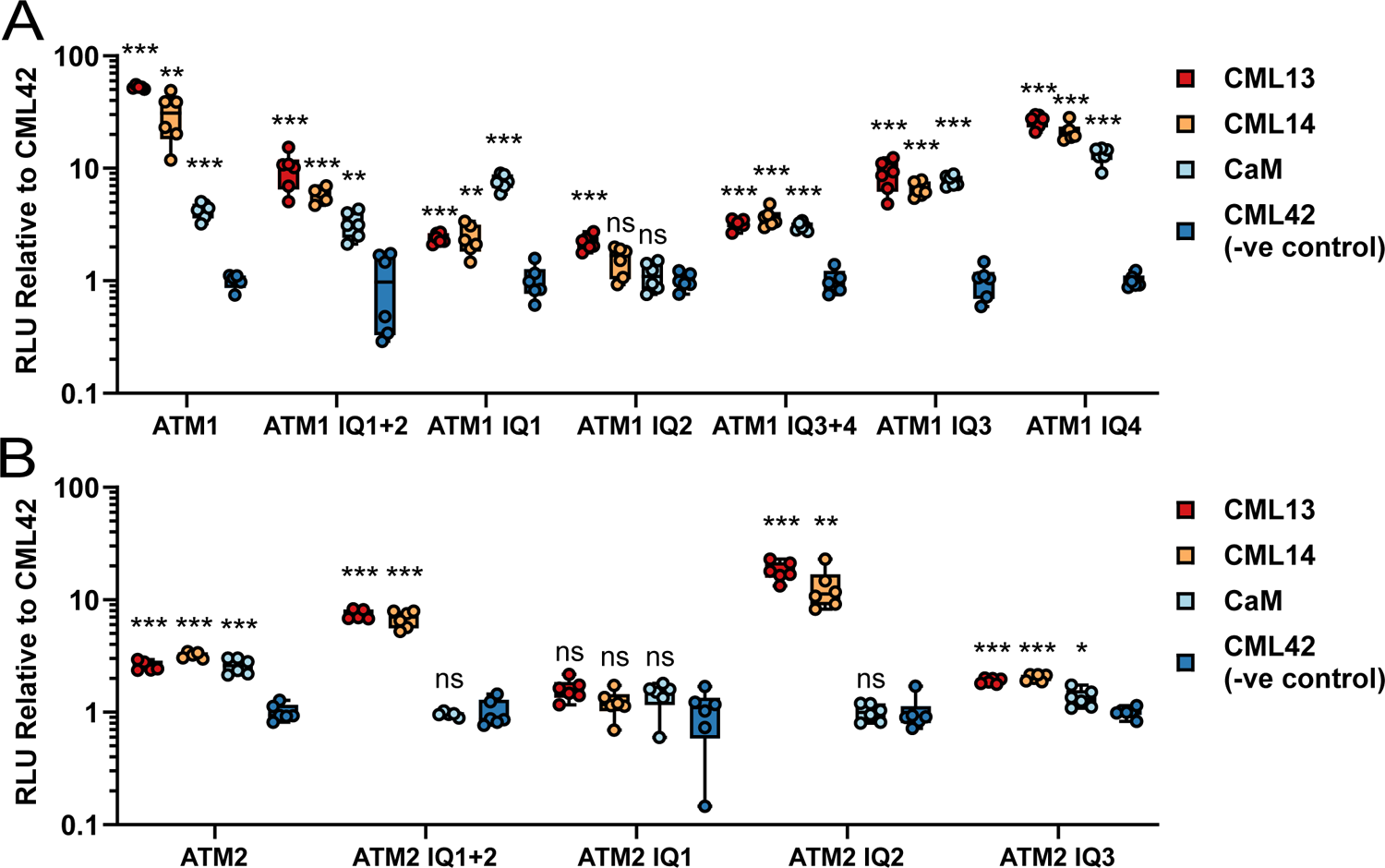
Split-luciferase protein interaction of Arabidopsis CaM, CML13, and CML14 with single and paired IQ-domains of myosin ATM1 and ATM2 *in planta*. *N. benthamiana* leaves were infiltrated with *Agrobacterium* harboring respective NLuc-prey (myosin VIII IQ domains) and CLuc-bait (CaM, CMLs) vectors and tested for luciferase activity 4 days later, as described in Materials and Methods. (**A**) Split-luciferase data are expressed as a fold-increase or -decrease relative to the RLU signal (log_10_ scale) observed using the negative control bait CML42 which was set to an RLU of 1. Boxes contain each data point for 6 technical replicates, means are shown by a horizontal bar, the colored region is the 95% confidence interval, and whiskers extend to maximum and minimum data points. Asterisks indicate a significantly higher signal vs CLuc-CML42 as a negative control bait (One-way ANOVA against CML42 with Sidak’s test for multiple comparisons, p-value * < 0.05, ** < 0.01, *** < 0.001). Data is representative of at least three independent experiments. RLU; relative light units.

Interestingly, for ATM2 binding assays, ATM2-IQ1+2 interacted with CML13 and CML14 but not with CaM (Fig. 4B). We did not test the combination of ATM2-IQ2+3 as these IQ domains are not arranged in close-proximity and are thus are not classified as a pair (Fig. 1A). Delineation of the individual IQ domains of ATM2, revealed that CML13 and CML14, but not CaM, interacted significantly with ATM2-IQ2, whereas the ATM2-IQ1 domain alone did not associate with any of the putative MLCs tested (Fig. 4B). ATM2-IQ3 bound to each of CaM, CML13, or CML14. In general, the delineation analysis of IQ domains within the neck regions of ATM1 and ATM2 strongly suggests specificity for different MLCs among some of the myosin VIII IQ motifs.

### Most myosin VIII tandem-IQ motifs have calcium-insensitive interactions with MLCs

We tested the effects of Ca^2+^ on the binding of putative MLCs to ATM1 and ATM2 using *in vitro* protein-interaction overlay assays (Fig. 5). As previously reported for other myosins, we observed poor solubility of recombinant fusion proteins that contained multiple IQ domains, and thus used myosin-fusion proteins solubilized in urea to testing binding to the full neck regions of ATM1 and ATM2 (Teresinski *et al*., 2023). Shorter regions of the neck domains were soluble as fusion proteins or synthetic peptides. We spotted 200 ng of pure myosin-fusion proteins onto nitrocellulose and probed these for interaction with fluorescently labeled CaM, CML13, or CML14 at a final concentration of 200 nM in the presence of excess Ca^2+^ or EGTA (apo-CaM/CML). Coomassie-stained blots in the uppermost panels indicate equivalent protein loading in all overlay assays. We did not observe the binding of any of these labeled proteins to a negative control, GB1, which was used as a fusion protein (pET21b-GB1). Each of these putative MLCs interacted with the tandem IQ domains of ATM1 and ATM2 independently of Ca^2+^ although qualitative signals were typically stronger when Ca^2+^ was present in the binding assays. However, we did observe some exceptions to this pattern. For example, the fluorescent signal of CML14 interaction with the full neck region of ATM2 was comparable in either the presence or absence of Ca^2+^. Interestingly, in contrast to our observations with the *in planta* SL assay (Fig. 4), CaM interacted with the ATM2 IQ1+2 peptide in overlay assays. Whether this reflects a difference in cellular vs *in vitro* conditions is unclear at this point. As an additional control, we labeled CML42 and tested its interaction with the full neck region of ATM1 and ATM2. We observed a faint but discernible signal only for CML42 with ATM2 under Ca^2+^ conditions, and these signals were notably weaker than those observed using CaM, CML13, or CML14 with ATM2, validating our use of CML42 as a negative control.

**Figure 5.**
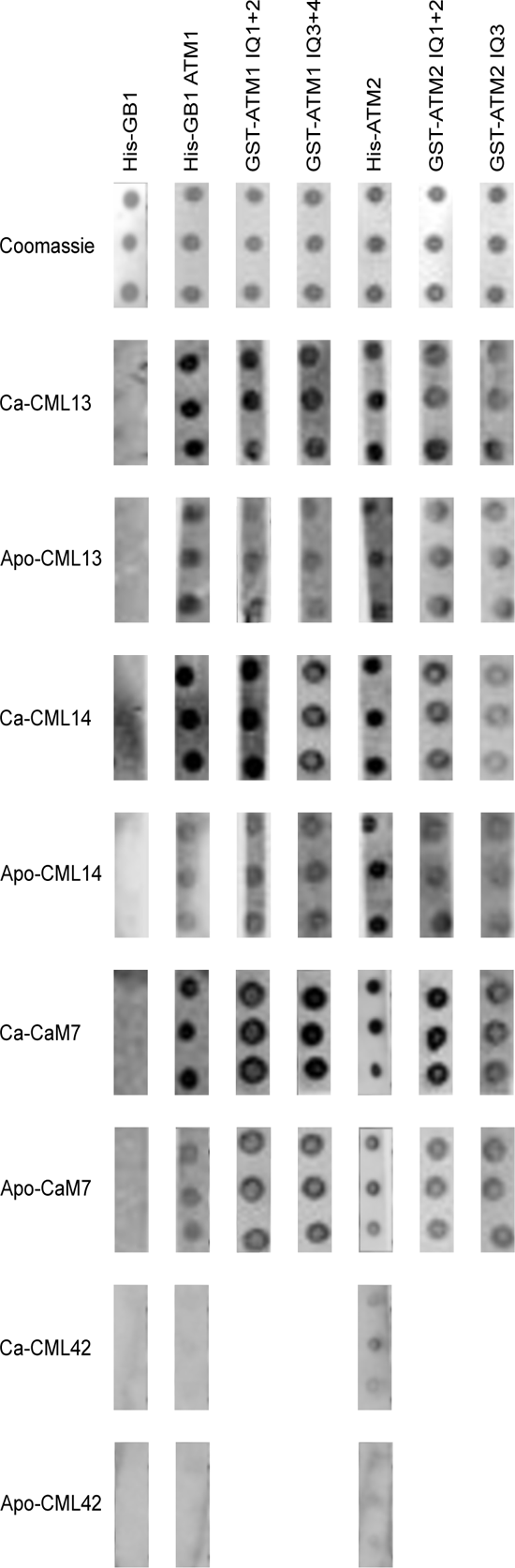
*In vitro* protein-interaction overlay assays of CaM, CML13, CML14, and CML42 with IQ domains of ATM1 and ATM2. Triplicate samples (200 ng) of pure, recombinant proteins ATM1 or ATM2 full-neck region (ATM1, ATM2), paired IQ domains (ATM1-IQ1+2, -IQ3+4, ATM2-IQ1+2), or the isolated IQ domain of ATM2 (ATM2-IQ3), were spotted onto nitrocellulose, blocked with 5% casein in TBST, and incubated with 200 nM of CaM, CML13, CML14, or CML42, as indicated, each of which was covalently labeled with the infra-red dye, 680RD-NHS as described in Materials and Methods. Recombinant GB1 protein (pET28-GB1) was tested as a negative control. Protein-protein interaction was assayed in the presence of 2 mM CaCl_2_ (Ca) or 5 mM EGTA (Apo) and detected using the LI-COR Odyssey-XF infra-red imager. Representative Coomassie-stained blots are presented along the top row. Data are representative of a minimum of three independent experiments. See Supplementary Table 2 for a description of the primary sequence from the neck regions of ATM1 and ATM2 that were tested for binding.

As an additional test of *in vitro* interaction, we labeled CaM, CML13, and CML14 with dansyl chloride and examined their binding in solution to synthetic peptides corresponding to the single IQ domains, ATM-IQ1 or ATM2-IQ1. Figure 6 shows the fluorescence spectra of dansylated (D-) CaM, -CML13, and -CML14 in the presence or absence of ATM1-IQ1 and ATM2-IQ1 peptides under Ca^2+^ vs EGTA conditions. In agreement with our *in planta* SL and *in vitro* overlay assays, ATM1-IQ1 associated with CML13, CML14, and CaM in both the presence and absence of Ca^2+^, as seen by the blue peak shift and increase in fluorescence intensity in the emission spectra (Fig. 6A).

**Figure 6.**
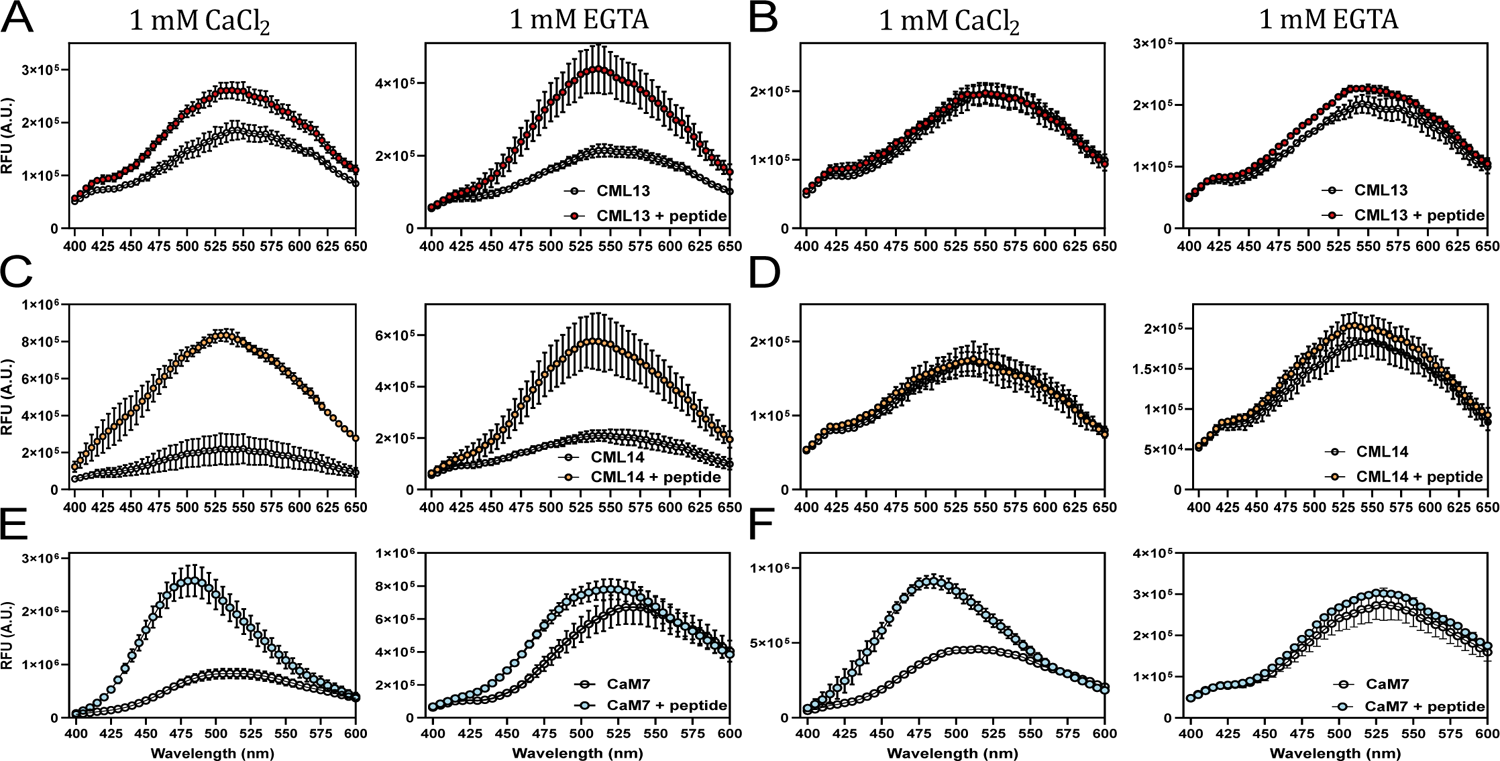
*In vitro* interaction of dansyl-CaM, -CML13, or -CML14 with IQ-domain synthetic peptides of ATM1-IQ1 and ATM2-IQ1. Dansyl fluorescence was measured over an emission wavelength window from 400 to 600 or 650 nm and an excitation wavelength of 360 nm. Samples of 3 µM dansyl-CML13, -CML14, or using 600 nM dansyl-CaM, were separately tested for fluorescence alone, or in the presence (A, C, E, respectively) of ATM1-IQ1, or (B, D, E, respectively) ATM2-IQ1 peptide under conditions of Ca^2+^ (left panels) or EGTA (right panels). Peptide concentrations were used at a 10-fold molar excess. Spectra were collected for dansyl-CaM and -CMLs in the presence or absence of IQ-peptides. The intensity was measured in arbitrary relative fluorescence units (RFU).

Interestingly, for ATM2-IQ1, a change in the spectrum of D-CaM was observed in the presence of 1 mM CaCl_2_ but not in EGTA, and the presence of ATM2-IQ1 did not alter the spectra of D-CML13 and D-CML14 (Fig. 6B). The apparent lack of interaction of CML13 and CML14 with ATM2-IQ1 here is consistent with our observations using the SL system (Fig. 4B). The effect of Ca^2+^ vs EGTA was most apparent in the case of CaM in the presence of either ATM1-IQ1 (Fig. 6E) or ATM2-IQ1 peptide (Fig. 6F), where Ca^2+^ elicited a marked increase in dansyl fluorescence and a blue shift of the emission spectrum.

### CML13, CML14, and CaM act as MLCs for myosin VIIIs motility

We generated two constructs to measure the velocities of ATM1 and ATM2. The ATM1 construct encodes a protein with the motor domain and native neck regions (four IQ motifs) of ATM1. Similarly, the ATM2 construct encodes the motor domain and native neck regions (three IQ motifs) of ATM2. These constructs were expressed using a baculovirus system in *High Five* insect cells and purified by affinity chromatography. The velocities of ATM1 and ATM2 were measured using an anti-myc antibody-based version of the *in vitro* actin gliding assay (Ito *et al*. 2007). The velocities of ATM1 and ATM2 were tested in the presence of CaM, CML13, and CML14 alone or in different combinations of two putative MLCs (Fig. 7A and B). The velocities of ATM1 with any single MLC and two different CMLs had the same activity in vitro (Fig. 7A). However, the combination of CaM and CML13 or CML14 showed increased motility and the fastest motility was observed using the combination of CaM with CML14 (Fig. 7A). In contrast, the velocity of ATM2 with either CML13 or CML14 alone was significantly slower compared to assays using CaM as the solo MLC. This is consistent with our interaction data that suggests that neither CML13 nor CML14 interact with ATM2-IQ1 (Figs. 4B, 6B, D, F). Similar to ATM1 assays, the motility of ATM2, was fastest when tested using combinations of CaM and CML13 or CML14 (Fig. 7B).

**Figure 7.**
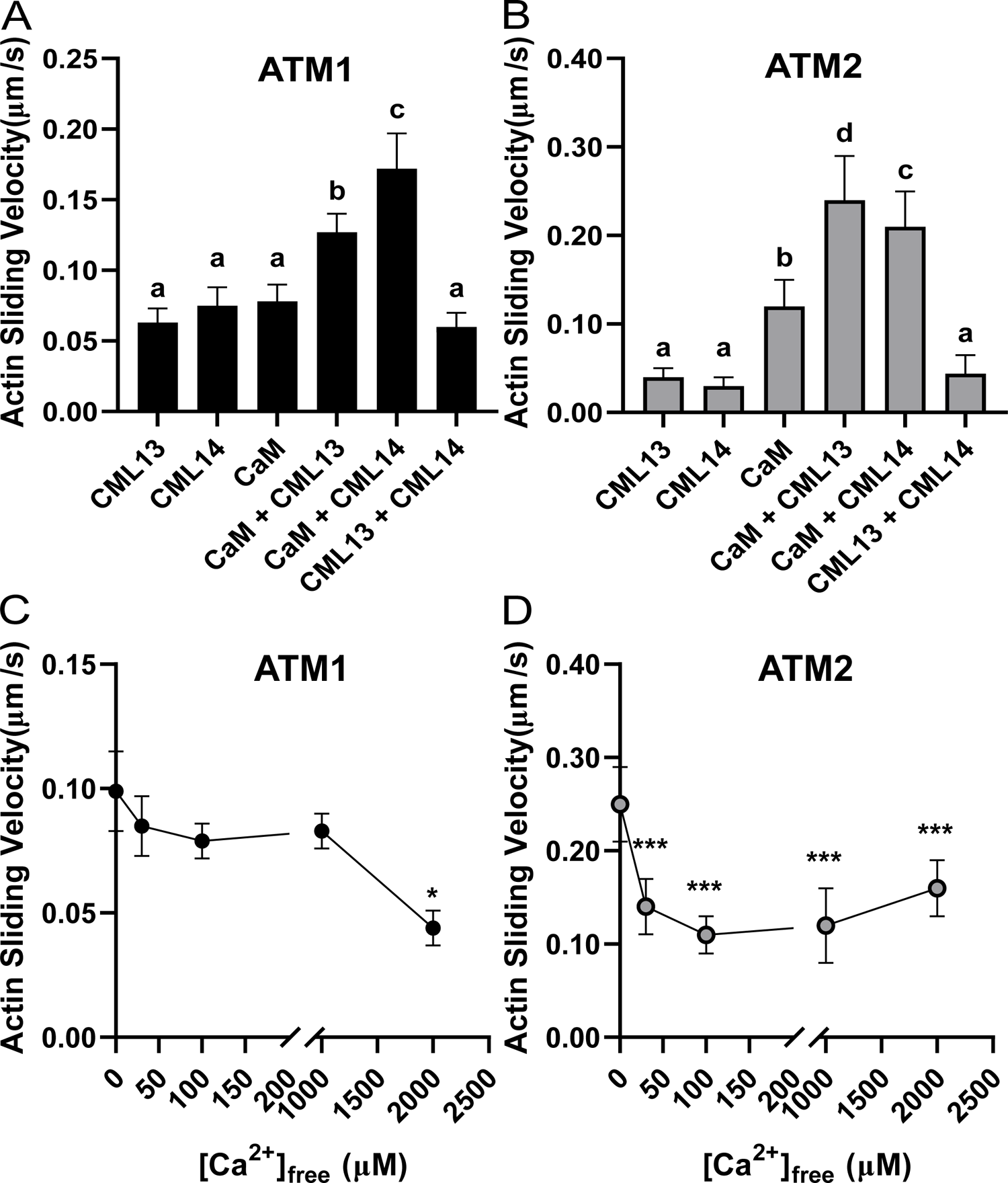
Actin-sliding assays indicate that CaM, CML13, and CML14 can function as light chains for ATM1 and ATM2. **A)** Actin sliding velocity of ATM1 in the presence of CaM, CML13, and CML14 alone or in different combinations of two putative MLCs. The concentrations used for CaM, CML13, and CML14 were 10µM, 30µM, and 30µM, respectively. **B)** Actin sliding velocity of ATM2 in the presence of CaM, CML13, and CML14 alone or in different combinations of two putative MLCs. The concentrations used for CaM, CML13, and CML14 were each 30 µM. **C)** The Ca^2+^ sensitivity of ATM1 in the presence of CaM and CML13. The concentrations used for CaM and CML13 were 10 µM and 30 µM, respectively. **D.** The Ca^2+^ sensitivity of ATM2 in the presence of CaM and CML13. The concentrations used for CaM and CML13 were 10 µM and 30 µM, respectively.

**Figure 8.**
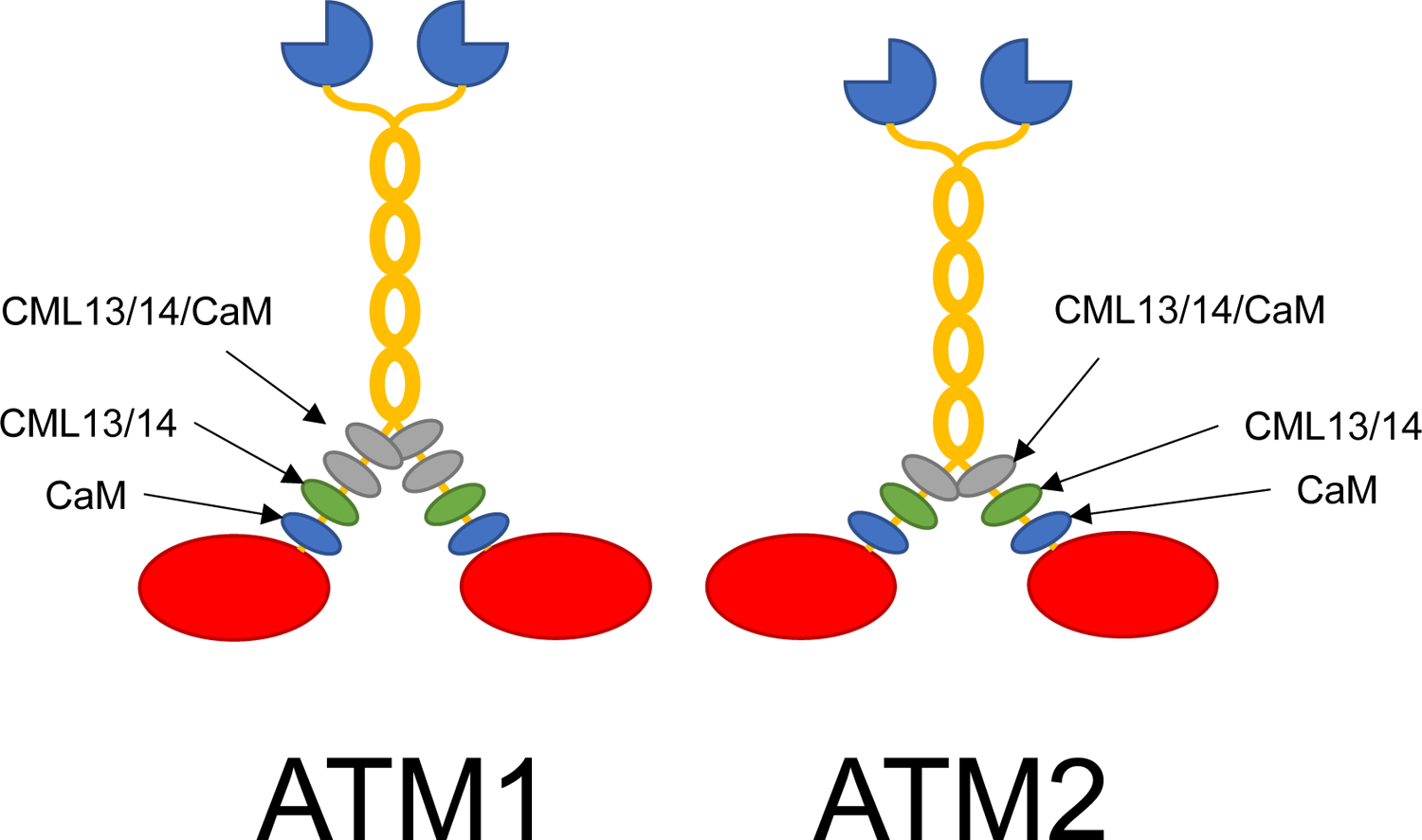
Working model of CaM, CML13, and CML14 as light chains for Arabidopsis myosins ATM1 and ATM2. Actin-binding, catalytic myosin head domains are presented in red, the neck regions are decorated with light chains represented as blue (CaM), green (CML13/14), or grey (any of CaM/CML13/14) ovals, the coiled-coil dimerization regions are in yellow, and the cargo-binding tail domains as dark blue, three-quarter circles. Light chains bind to the IQ domains within the neck region of myosins to provide the leverage and structural integrity needed for the power stroke. We speculate that CaM is the preferred light chain at the IQ1 “essential” position (nearest the head) for both ATM1 and ATM2 whereas CML13 or CML14 are preferred light chains at IQ2, in the “regulatory” position. Other IQ domains did not show a clear preference in our tests and may be occupied by CaM, CML13, CML14, or possibly other light chains.

The Ca^2+^ sensitivity of ATM1 and ATM2 was tested using CaM and CML13 as representative MLCs under conditions of increasing concentrations of free Ca^2+^ (Fig. 7C and D). ATM1 displayed reduced motility only at high Ca^2+^ concentrations (i.e. above 1 mM), whereas ATM2 activity was inhibited by much lower Ca^2+^ concentrations, with a marked reduction in motility observed at 30 µM free Ca^2+^ (Fig. 7D). Taken together, our actin motility assays clearly indicate that along with CaM, both CML13 and CML14 are able to function as MLCs.

## DISCUSSION

Despite the importance of MLCs for myosin function, relatively little is known about the identity and properties of plant MLCs. While previous reports have indicated that CaM likely functions as an MLC in plants (Yokota and Shimmen, 1994; Haraguchi *et al*., 2014), evidence to support speculation that plants also use non-CaM MLCs has been limited (Haraguchi *et al*., 2018). Multiple independent lines of analysis in our study support our hypothesis that Arabidopsis CML13 and CML14 function as MLCs given that: (i) they bind specifically to all four myosin VIIIs *in planta* (Figs. 2, 4), (ii) they interact at physiological concentrations (i.e. 200 nM) with IQ domains from myosin VIIIs *in vitro* (Fig. 5), (iii) they co-localize with myosin VIIIs when co-expressed in plant cells (Fig. 3), and (iv) they support *in vitro* myosin activity (Fig. 7).

The tandem IQ domains of a given myosin allow multiple MLCs to bind and confer structural integrity and, in some cases, impact activity (Heissler and Sellers, 2014). Different MLCs often possess specificity for distinct IQ domains within a given myosin, where one MLC occupies the first IQ position and is designated the ELC and a different MLC occupies the second IQ as the RLC (Heissler and Sellers, 2014). By convention, additional light chains, binding to other IQ domains toward the myosin tail, are generally not referred to as essential or regulatory. However, the designations of CaM or specific CaM-related proteins as strictly an ELC or RLC for a given myosin should be approached cautiously given the complexity of *in vivo* associations (Heissler and Sellers, 2014). For example, a specific MLC that binds to the first (ELC position) or second (RLC position) IQ of a particular myosin, does not exclude it from serving as an MLC at other IQ domains within that myosin. Although the IQ motif is broadly defined according to the IQXXXRGXXXR consensus, there is considerable variation among IQ domains (Bähler and Rhoads, 2002; Vetter and Leclerc, 2003).

The canonical CaM-binding IQ motif has a large hydrophobic residue immediately upstream of the first consensus Arg residue and another following the conserved Gly and second Arg: IQXXΦRGΦXXRXXΦ, where Φ refers to large hydrophobic residues (Houdusse *et al*., 2006). It is noteworthy that the first IQ domain (IQ1) for all four Arabidopsis myosin VIII isoforms adheres to this pattern (Fig. 1A). In contrast, although the IQ2 domains of these myosin VIIIs are highly conserved, they possess smaller hydrophobic residues (e.g. Val, Iso) preceding the Arg-Gly consensus, followed by a Glu (Fig. 1A). Negatively-charged residues within IQ domains, and other CaM-binding domains, have been reported to impede interaction with CaM (Vetter and Leclerc, 2003; Slaughter *et al*., 2005). These notable differences in primary sequence suggest that the IQ1 and IQ2 domains of Arabidopsis myosin VIIIs may be occupied by non-CaM MLCs, consistent with our observations. Structurally, how the sequence variation among these IQ domains and the peripheral regions impacts MLC specificity, binding affinity, and myosin VIII function remains an open question.

Using the SL *in planta* protein-interaction system we observed some differences in MLC preference among the IQ motifs of ATM1 and ATM2 as representative myosin VIII isoforms (Fig. 4). Despite the divergence of ATM1-IQ3 from the canonical IQ consensus motif, it bound CaM, CML13, and CML14 with comparable signal strength (Fig. 4A), indicating that all essential residues for interaction with different MLCs are present. Given the adherence of ATM1-IQ4, and ATM2-IQ3 to the IQ consensus, it is not surprising that they bound CaM, but this data also indicates that CML13 and CML14 can bind to both conserved and divergent IQ domains (Fig. 4A, B). In contrast, we observed specificity of ATM2-IQ2 with CML13 and CML14, and ATM1-IQ2 with CML13 alone (Fig. 4A, B), suggesting that CML13 and CML14 likely act as the RLCs for these myosins. We speculate that the conserved Glu at IQ domain position 8 within the consensus motif of IQ2 in myosin VIIIs may decrease CaM affinity without impacting CML13/14-IQ interaction. Furthermore, the specificity differences between ATM1-IQ2 and ATM2-IQ2 may result from the Lys/Arg residues at position 9 and/or the Glu/Leu residues at position 13, respectively (Houdusse *et al*., 2006). However, analysis of solved structures will be needed in the future to fully elucidate the binding properties of IQ/CML complexes.

Collectively, our study indicates that CaM, CML13, and CML14 interact with myosin VIII IQ domains, but it should be pointed out that our data from *in planta* and *in vitro* assays did not align in all cases, a phenomenon previously reported for analyses of CaM binding to IQ domains in plant proteins (Bürstenbinder *et al*., 2013). For example, ATM2-IQ1 did not interact with any of the MLCs tested in the SL assay (Fig. 4B), whereas CaM showed strong interaction with GST-ATM2-IQ1+2 and an ATM2-IQ1 synthetic peptide *in vitro* as judged by overlay assays (Fig. 5) or dansyl-CaM binding spectrophotometry (Fig. 6), respectively. These differences may reflect the cellular conditions or sensitivity limitations of the SL system, or the simplicity of the *in vitro* assays where there is an absence of CaM/CML binding competitors. Regardless, as CaM alone was able to support ATM2 motility to a greater level than either CML13 or CML14 alone (Fig. 7), this represents strong evidence that it can serve as an ELC for ATM2. We are unaware of any previous *in vitro* studies on the motility of ATM2 or the identity of its MLCs. Thus, when considered together with our binding and motility data (Fig. 5, 6, 7), the collective evidence suggests that CaM is likely an ELC for ATM1, ATM2, and probably myosin VIIIs in general.

It is also interesting that we observed weak signals for ATM1-IQ2 interaction with each of the MLCs tested using the *in planta* SL system, with only CML13 emerging as a putative interactor (Fig. 4). Despite this, *in vitro* motility of ATM1 was supported by a combination of CaM and either CML13 or CML14 (Fig. 7A). In actin-sliding (motility) assays, it is thought that all IQ domains need to be occupied for smooth actin movement (Haraguchi *et al*., 2018). Given that we observed such smooth ATM1 activity with CaM and CML14 even in the absence of CML13, this suggests that they can function as MLCs at ATM1-IQ2, at least under *in vitro* conditions. However, as CML13, but not CML14 or CaM, interacted with ATM1-IQ2 in the *in planta* SL interaction assays (Fig. 4), this raises the question as to whether CML13 might be the preferred RLC *in vivo*. Although we used the well-established *High Five* insect cell expression system (Vaughn *et al*., 1977; Haraguchi *et al*., 2014) to prepare myosin for activity assays, one caveat of this method is that we cannot exclude the possibility that other proteins, including conserved CaM, copurified with our preparations. However, as CMLs are unique to plants, and their presence increased ATM1 and ATM2 *in vitro* activity, our data suggest that *High Five* cells do not possess proteins that can substitute for CML13 or CML14 as MLCs (Haraguchi *et al*., 2018).

We also corroborated earlier work that suggested CML24 is a putative interactor with some myosin VIII isoforms (Abu-Abied *et al*., 2006). CML24 expression is upregulated in response to mechanical stimulation and is thought to be important in thigmomorphogenesis in Arabidopsis (Wang *et al*., 2011; Darwish *et al*., 2022). Furthermore, *cml24* point mutants display actin and cortical microtubule defects (Wang *et al*., 201). These earlier studies led to speculation that CML24 may interact with myosins during mechanical stress to aid in the organization of the cytoskeleton (Abu-Abied *et al*., 2006; Wang *et al*., 2011). Although we showed the *in planta* interactions of CML24 with the neck domains of several myosin VIII isoforms (Fig. 2C), it generally exhibited weaker SL signals than CaM, CML13, or CML14. Thus, we restricted our co-localization, *in vitro* binding, and myosin motility assays to the latter group. As such, further studies will be needed to assess whether CML24 serves as a *bona fide* MLC.

Myosin VIIIs have been suggested to function in the organization of the cytoskeleton at the plasma membrane (Golomb *et al*., 2008; Bar-Sinai *et al*., 2022). The punctate localization pattern that we observed (Fig. 3) using myosin VIIIs comprised of neck and tail regions is consistent with earlier studies and is thought to reflect the location of myosin-VIII cargo to which the tail domains bind (Golomb *et al*., 2008). Our colocalization data indicate that myosin VIIIs recruit CML13 and CML14 as MLCs to the plasma membrane as components of the acto-myosin cytoskeleton (Fig. 3). Myosin VIII-A and VIII-B typically displayed a more diffuse localization compared to ATM1 and ATM2, consistent with previous reports (Avisar et al., 2009). Interestingly, Pearson’s coefficients of colocalization for CML13 and CML14 were lowest with myosin VIII-B, but whether this indicates a weaker binding affinity for this isoform remains unknown. Although we did not test all 50 Arabidopsis CMLs, our SL analysis indicates clear specificity among the CML family for myosin VIII interaction (Fig. 2). However, this does not exclude the possibility that other CMLs or non-CML proteins might also function as MLCs. It is also noteworthy that while CaM showed clear colocalization with myosin VIIIs, Pearson’s coefficient values were generally lower than those for CML13 and CML14 for each respective myosin other than VIII-B. Various CaM-binding proteins in these cells may be competing for GFP-CaM in these assays, reducing its availability for myosins, but this remains speculative. Although the weakest correlation of colocalization was between ATM1 and CaM, our SL analyses, *in vitro* binding assays, and motility tests all demonstrate that CaM can function as an MLC with ATM1, supporting a previous report (Haraguchi *et al*., 2014).

Myosin VIIIs associate with the actin network and localize to plasma membranes, plasmodesmata, endosomes, and other internal structures, suggesting a breadth of roles within cells (Golomb *et al*., 2008; Haraguchi *et al*., 2014; Nebenführ and Dixit, 2018; Bar-Sinai *et al*., 2022). However, insight into their functions has been hindered by an apparent genetic redundancy among myosins. Although no obvious phenotype was initially observed in an Arabidopsis mutant lacking all four myosin VIIIs (Talts *et al*., 2016), a subsequent study reported a subtle phenotype for *atm1* single knockouts, a reduced hypocotyl elongation that was recoverable by sucrose supplementation (Olatunji and Kelley, 2020). Recently, ATM1 was implicated in root apical meristem organization (Olatunji *et al*., 2022) and myosin VIIIs also appear to function in *Agrobacterial* transformation of Arabidopsis cells (Liu et al., 2023). The use of *Physcomitrella patens* as a model demonstrated that the loss of all five myosin VIIIs resulted in a pleiotropic phenotype that included reduced size and growth rate of gametophytes, suggesting roles in cell expansion and hormone homeostasis (Wu *et al*., 2011).

In general, our data provide strong evidence that CaM, CML13, and CML14 are *bona fide* Arabidopsis MLCs. The importance of Ca^2+^ in this interaction is unclear and may vary depending on cellular conditions. In the yeast or mammalian class V myosin model, CaM is thought to be the RLC and is responsible for the Ca^2+^ regulation of the myosin V holoenzyme by dissociating from IQ2 in the presence of Ca^2+^ (Batters and Veigel, 2016). However, the physiological relevance of this Ca^2+^-induced dissociation remains unclear, given that under *in vitro* conditions 100 µM levels or more of Ca^2+^ are often required to dissociate CaM from the IQ domains (Yokata *et al*., 1999; Manceva *et al*., 2007). The *in vivo* picture is thus likely quite complex, and the role of Ca^2+^ may vary among MLCs and specific IQ domains within a given myosin.

Similar to reports on other, unconventional myosins, such as class Vs and XIs (Tominaga and Nakano, 2012; Batters and Veigel, 2016), myosin VIIIs in our assays behaved as Ca^2+^-independent but not completely Ca^2+^-insensitive motors, where ATM1 and ATM2 motor-neck holoenzymes exhibited decreasing actin sliding velocities in the presence of MLCs and increasing Ca^2+^ concentrations (Fig. 7). However, this model does not fully explain the broader plant myosin data in the literature, and it has been speculated that the myosin V paradigm is not applicable to plant myosins (Batters and Veigel, 2016). Unlike the myosin V model, the disassociation of CaM from the lower affinity IQ2 is not likely how myosin VIIIs are regulated by Ca^2+^. Instead, as CML13 and CML14 are mainly Ca^2+^ insensitive (Vallone *et al*., 2016; Teresinski *et al*., 2023), one possibility is that a conformational change or an increase in the rate of dissociation of Ca^2+^-CaM at IQ1 regulates Ca^2+^ sensitivity as has been reported previously for myosin Va and Ic, respectively (Manceva *et al*., 2007; Shen *et al*., 2016). Our *in vitro* binding data (Fig. 6E, F), showing a Ca^2+^-induced conformation change in CaM when bound to ATM1-IQ1 and ATM2-IQ1, is consistent with this speculation. An alternative hypothesis considers the impact of competition for CaM between myosins and other CaM-binding proteins under conditions of elevated Ca^2+^. CML13, CML14, and possibly other MLCs may occupy some IQ domains of plant myosins and afford them structural stability when Ca^2+^ levels rise in response to stimuli and free CaM levels are limiting due to CaM interaction with various targets. Indeed, our discovery of CML13/14 as novel MLCs may help explain discrepancies between previous models and observations. For example, when Arabidopsis myosin XIs were tested with CaM as the sole MLC, several isoforms lacked motility (Haraguchi *et al*., 2018), likely due to the absence of other MLCs, such as CML13 and/or CML14.

Another area of future research will be to extend interaction and motility assays of CML13 and CML14 to the larger myosin XI family, comprised of 13 members. Like myosin VIIIs, myosin XIs are only found in the green lineage and have been linked to roles in cytoplasmic streaming, cell expansion, trichome branching, fertility, as well the shape and movement of organelles and vesicles (Nebenführ and Dixit, 2018). A major challenge in elucidating specific roles for CML13 and CML14 concerns their potential number of targets, including CAMTAs, IQD proteins, and myosins (Teresinski *et al*., 2023). In Arabidopsis, these putative targets represent 56 different proteins, each with multiple IQ domains, and thus, like CaM, the number and variety of cellular processes that CML13/14 may participate in is expansive.

Speculation about the presence of non-CaM MLCs in plants was raised several decades ago but, to our knowledge, the present study is the first to empirically demonstrate that specific CMLs can function as MLCs (Vahey *et al*., 1982; Ma and Yen, 1989). The identification of CML13 and CML14 as novel MLCs should help accelerate myosin research. Among the key, unresolved questions to be addressed are whether CML13/14 function as class XI MLCs, how the specificity of CaM or CML-IQ interaction is achieved, and whether the phosphorylation of CML13 and CML14 impacts myosin activity/function.

## Acknowledgements

We thank Dr. Zongchao Jia (Queen’s University) for the kind gift of plasmid pET28b-GB1.

## Author Contributions

WS, KS, KI, ES, MT conceived the experimental design, KS, BH, HJT, VD, EB, S B-S, and TH performed experiments and collected data, and all authors contributed to data interpretation and manuscript writing.

## Data Availability

All data supporting the findings of this study are available within the paper and its supplementary materials published online.

## Supplementary Materials

### Supplementary Material Legends (brief)

**Supplementary Table T1.**
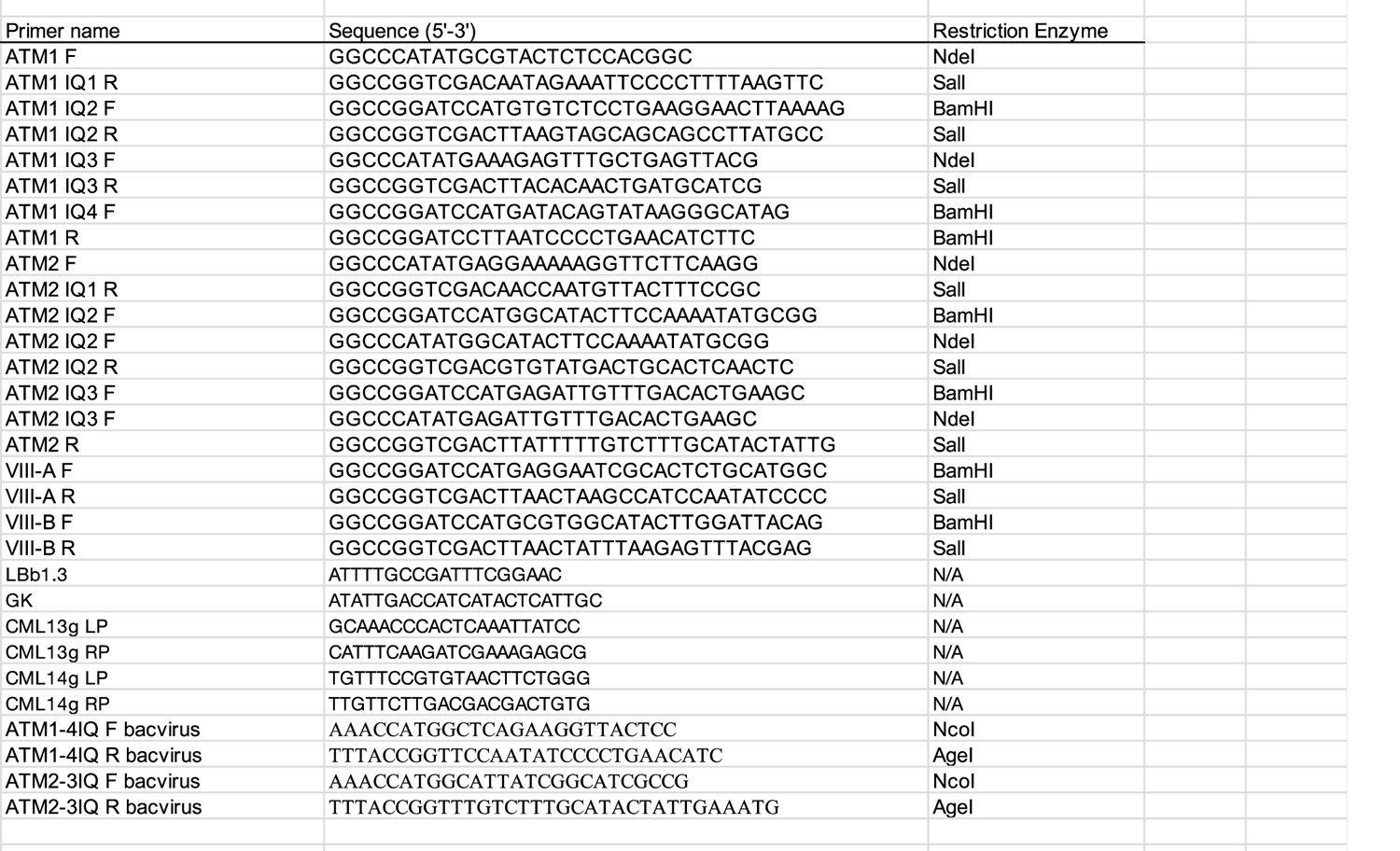
List of oligonucleotide primers used in this study

**Supplementary Table T2.**
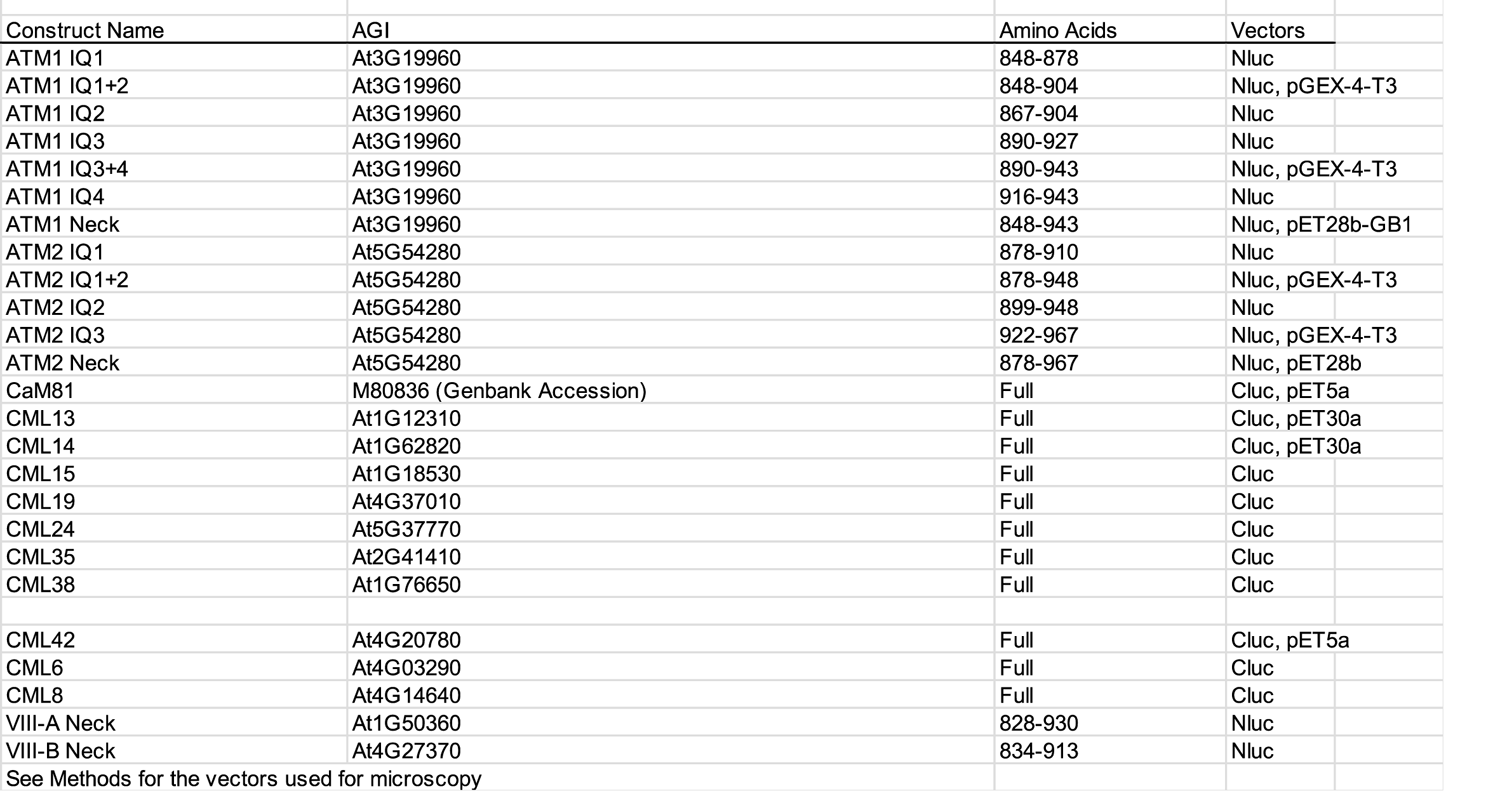
Description of plasmid constructs used in this study

**Supplemental Fig 1.**
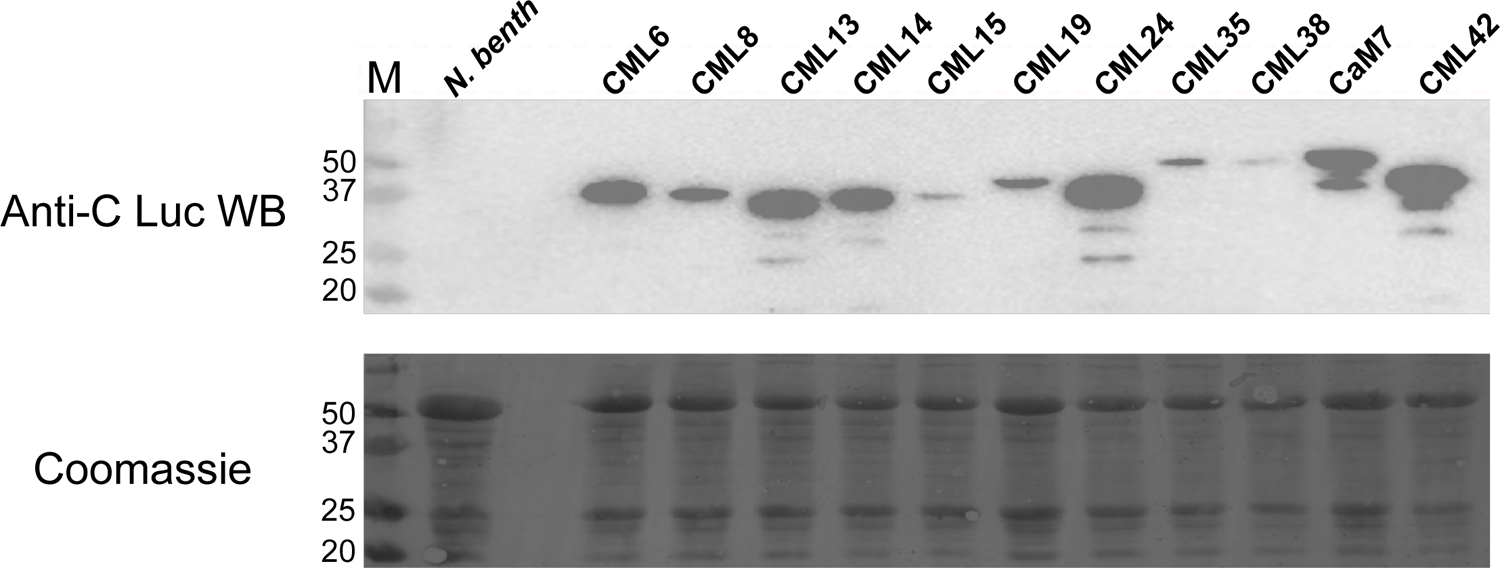
Immunoblot showing expression of C-Luc-CaM/CML fusion proteins in N. benthamiana. Leaves of N. benthamiana were infiltrated with Agrobacteria expressing C-Luc (bait) vectors and N=Luc (prey) vectors as described in Materials and Methods. Immunoblots (upper panel) using anti-C-terminal luciferase antisera, or Coomassie-stained gels showing equal lane loading, following SDS-PAGE of samples of total, clarified protein extracts (∼25ug) from leaves used in the split-luciferase assays.

**Supplemental Figure 2.**
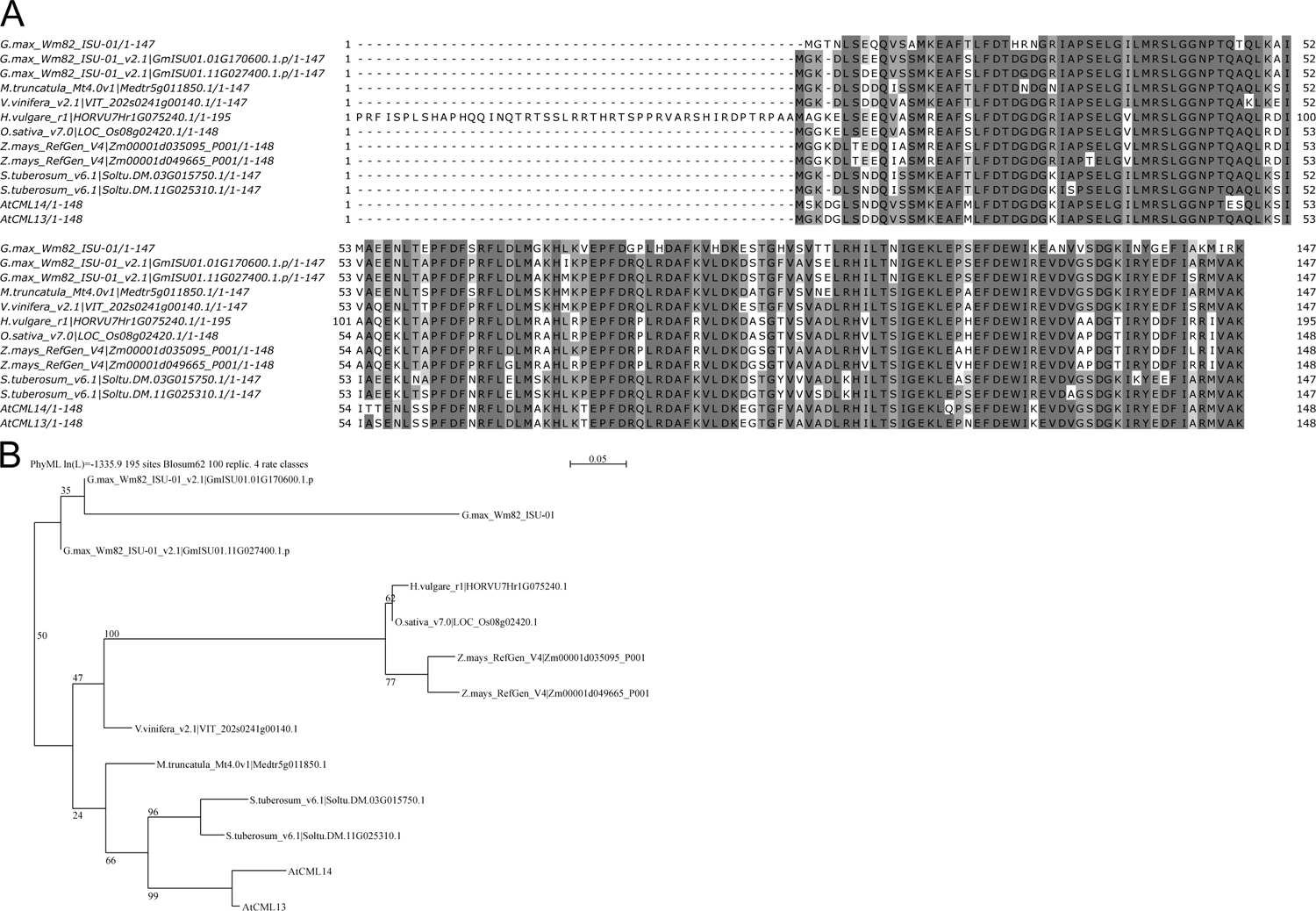
Comparison of Arabidopsis CML13 and CML14 with predicted orthologs. (A) Protein alignment of CML13 and CML14 with orthologs from various plant species. Amino acid residues were shaded based on their percent identity, dark grey if identical, and progressively lighter grey until white as unconserved. ClustalΩ was used for alignment (Sievers and Higgens, 2014) and images were generated using Jalview Version 2.11.2.6. (B) Phylogenetic tree showing relatedness among the proteinss compared in panel A. SeaView (v 5.0, Gouy et al., 2021) was used to generate the tree using neighbour-joining with bootstrapping analysis (1000 reiterations).

**Supplemental Fig 3.**
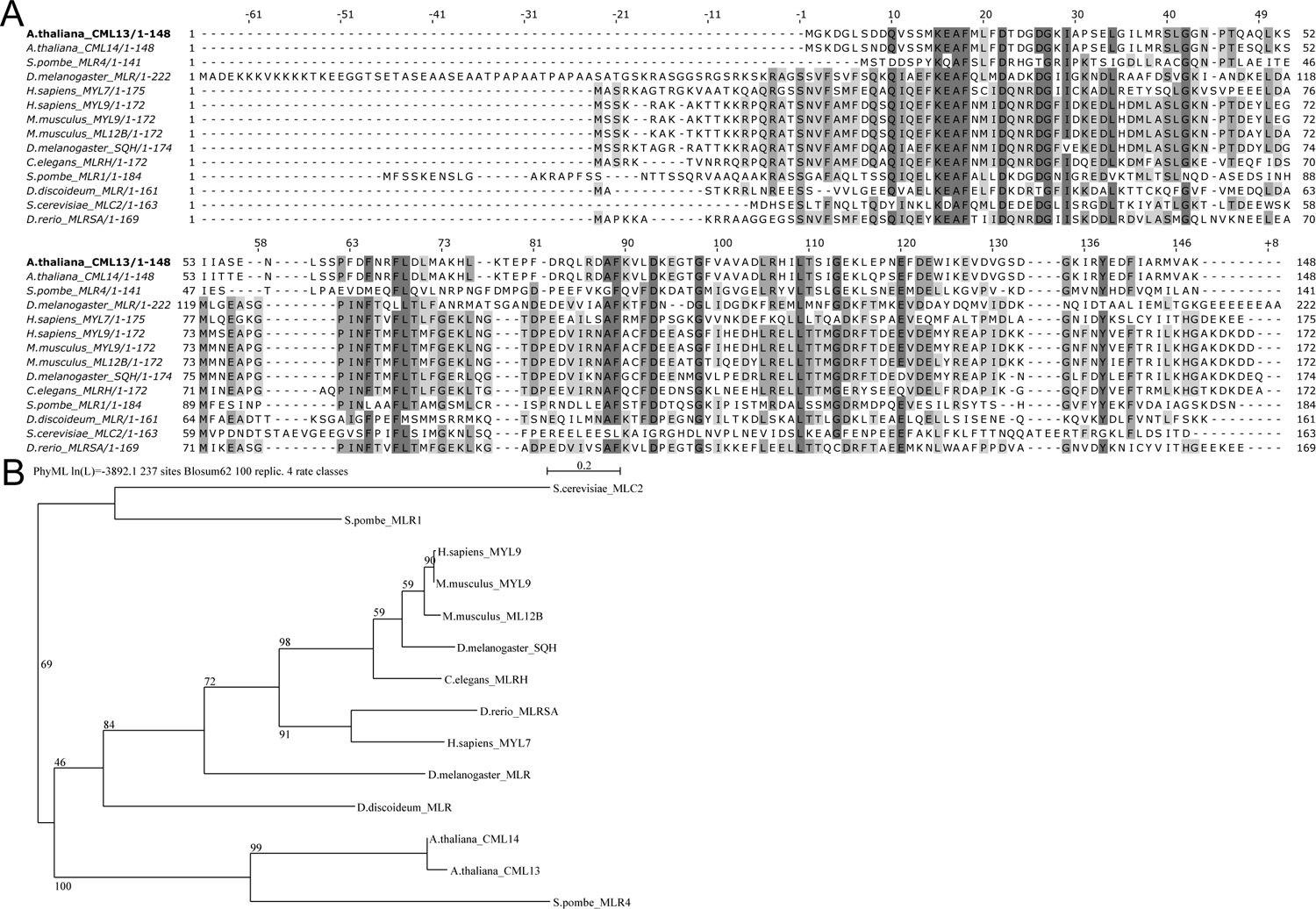
Comparison of Arabidopsis CML13/14 with regulatory myosin light-chains (RLCs) from various species of eukaryotes. (A) Protein alignment of CML13 and CML14 with myosin RLCs from multiple species. Amino acid residues were shaded based on their percent identity, dark grey if identical, and progressively lighter grey until white as unconserved. ClustalΩ was used for alignment (Sievers and Higgens, 2014) and images were generated using Jalview Version 2.11.2.6. (B) Phylogenetic tree showing relatedness among myosin RLCs compared in panel A. SeaView (v 5.0, Gouy et al., 2021) was used to generate the tree using neighbour-joining with bootstrapping analysis (1000 reiterations).

**Supplemental Figure 4.**
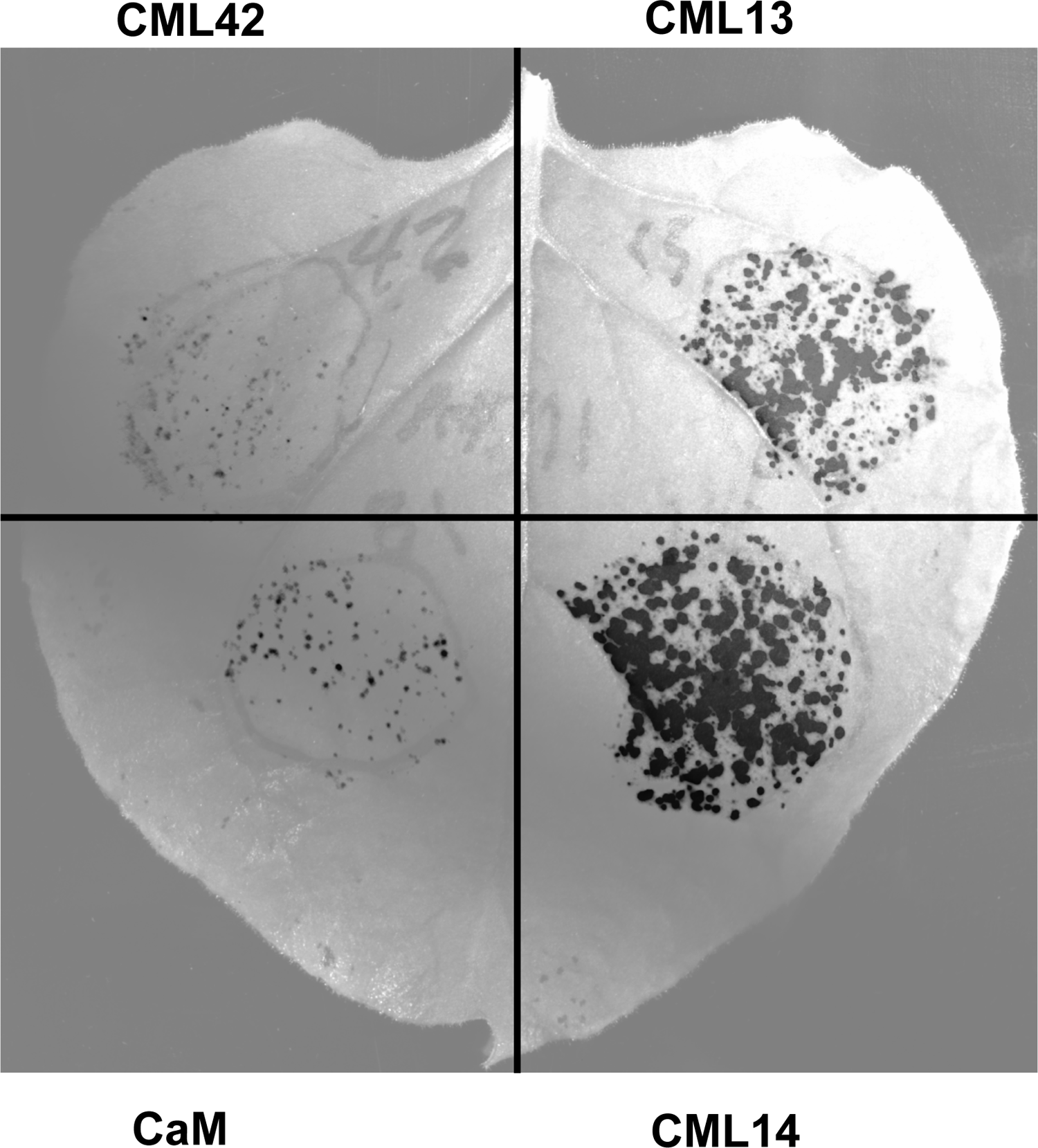
Whole-leaf image of split-luciferase protein-protein interaction bioluminescent assays using transiently transformed N. benthamiana. A representative image is shown for interaction analysis of CML13, CML14, CaM, and CML42 (negative control) as baits (as C-Luc fusion proteins) with the neck region of ATM1 (as N-luc fusion) prey protein.

## REFERENCES

Abu-Abied M, Belausov E, Hagay S, Peremyslov V, Dolja V, Sadot E. 2018. Myosin XI-K is involved in root organogenesis, polar auxin transport, and cell division. Journal of Experimental Botany 69, 2869–2881.

Abu-Abied M, Golomb L, Belausov E, Huang S, Geiger B, Kam Z, Staiger CJ, Sadot E. 2006. Identification of plant cytoskeleton-interacting proteins by screening for actin stress fiber association in mammalian fibroblasts. The Plant Journal 48, 367–379.

Alaimo A, Malo C, Areso P, Aloria K, Millet O, Villarroel A. 2013. The use of dansyl-calmodulin to study interactions with channels and other proteins. Methods in Molecular Biology 998, 217–231.

Altman, D. 2013. Myosin: Fundamental Properties and Structure. In: Roberts, GCK (eds) Encyclopaedia of Biophysics. Springer, Berlin, Heidelberg. https://doi.org/10.1007/978-3-642-16712-6_753

Avisar D, Abu-Abied M, Belausov E, Sadot E, Hawes C, Sparkes IA. (2009). A comparative study of the involvement of 17 Arabidopsis myosin family members on the motility of Golgi and other organelles. Plant Physiology 150, 700–709.

Bähler M, Rhoads A. 2002. Calmodulin signaling via the IQ motif. FEBS Letters 513, 107–113.

Baluška F, Cvrčková F, Kendrick-Jones J, Volkmann D. (2001). Sink plasmodesmata as gateways for phloem unloading. Myosin VIII and calreticulin as molecular determinants of sink strength? Plant Physiology 126, 39–46.

Bar-Sinai S, Belausov E, Dwivedi V, Sadot E. 2022. Collisions of cortical microtubules with membrane associated Myosin VIII tail. Cells 11, 145

Bashandy H, Jalkanen S, Teeri TH. 2015. Within leaf variation is the largest source of variation in agroinfiltration of *Nicotiana benthamiana*. Plant Methods 11, 47.

Batters C, Veigel C. 2016. Mechanics and activation of unconventional myosins. Traffic 17, 860– 871.

Belausov E, Dwivedi V, Yechezkel S, Bar-Sinai S, Sadot, E. (2023). Documentation of microtubule collisions with Myosin VIII ATM1 containing membrane-associated structures. *Methods in Molecular Biology* (Clifton, N.J.) 2604, 77–88.

Buchnik L, Abu-Abied M, Sadot, E. (2015). Role of plant myosins in motile organelles: is a direct interaction required? Journal of Integrative Plant Biology 57, 23–30.

Bürstenbinder K, Savchenko T, Müller J, Adamson AW, Stamm G, Kwong R, Zipp BJ, Dinesh DC, Abel S. 2013. Arabidopsis calmodulin-binding protein IQ67-domain 1 localizes to microtubules and interacts with kinesin light chain-related protein-1. The Journal of Biological Chemistry 288, 1871–1882.

Clapham DE. 2007. Calcium signaling. Cell 131, 1047–1058.

Darwish E, Ghosh R, Ontiveros-Cisneros A, Tran HC, Petersson M, De Milde L, Broda M, Goossens A, Van Moerkercke A, Khan K, Van Aken O. 2022. Touch signaling and thigmomorphogenesis are regulated by complementary CAMTA3- and JA-dependent pathways. Science Advances 8, eabm2091.

DeFalco TA, Bender KW, Snedden WA. 2009. Breaking the code: Ca^2+^ sensors in plant signaling. Biochemical Journal 425, 27–40.

DeFalco TA, Chiasson D, Munro K, Kaiser BN, Snedden WA. 2010. Characterization of GmCaMK1, a member of a soybean calmodulin-binding receptor-like kinase family. FEBS Letters 584, 4717–4724.

Duan Z, Tominaga M. (2018). Actin-myosin XI: an intracellular control network in plants. Biochemical and Biophysical Research Communications 506, 403–408.

Engler, C, Youles, M, Gruetzner, R, Ehnert, T-M, Werner, S, Jones, JDG, Patron, NJ, Marillonnet, S. (2014). A Golden Gate Modular Cloning Toolbox for Plants. ACS Synthetic Biology 3, 839–843.

Fischer C, DeFalco TA, Karia P, Snedden WA, Moeder W, Yoshioka K, Dietrich P. 2017. Calmodulin as a Ca^2+^-sensing subunit of Arabidopsis cyclic nucleotide-gated channel complexes. Plant and Cell Physiology 58, 1208–1221.

Goldstein, D. 1979. Calculation of the concentrations of free cations and cation-ligand complexes in solutions containing multiple divalent cations and ligands. Biophysical Journal 26, 235–242.

Golomb L, Abu-Abied M, Belausov E, Sadot E. 2008. Different subcellular localizations and functions of Arabidopsis myosin VIII. BMC Plant Biology 8, 3.

Gouy M, Tannier E, Comte N, Parsons DP. 2021. Seaview Version 5: A multiplatform software for multiple sequence alignment, molecularpPhylogenetic analyses, and tree reconciliation. Methods in Molecular Biology 2231, 241–260.

Haraguchi T, Ito K, Duan Z, Rula S, Takahashi K, Shibuya Y, Hagino N, Miyatake Y, Nakano A, Tominaga M. 2018. Functional diversity of class XI myosins in *Arabidopsis thaliana*. Plant and Cell Physiology 59, 2268–2277.

Haraguchi T, Tominaga M, Matsumoto R, Sato K, Nakano A, Yamamoto K, Ito K. 2014. Molecular characterization and subcellular localization of Arabidopsis class VIII myosin, ATM1. Journal of Biological Chemistry 289, 12343–12355.

Haraguchi T, Tamanaha M, Suzuki K, Yoshimura K, Imi T, Tominaga M, Sakayama H, Nishiyama T, Murata T, Ito K. 2022. Discovery of ultrafast myosin, its amino acid sequence, and structural features. *Proceedings of the National Academy of Sciences*, USA 119, e2120962119.

Hartman MA, Finan D, Sivaramakrishnan S, Spudich JA. 2011. Principles of unconventional myosin function and targeting. Annual Review of Cell and Developmental Biology 27, 133–155.

Heissler SM, Sellers JR. 2014. Myosin light chains: Teaching old dogs new tricks. Bioarchitecture 4, 169–188.

Henn, A, Sadot, E. (2014). The unique enzymatic and mechanistic properties of plant myosins. Current Opinion in Plant Biology 22, 65–70.

Houdusse A, Gaucher J-F, Krementsova E, Mui S, Trybus KM, Cohen C. 2006. Crystal structure of apo-calmodulin bound to the first two IQ motifs of myosin V reveals essential recognition features. *Proceedings of the National Academy of Sciences*, USA 103, 19326–19331.

Ito K, Ikebe M, Kashiyama T, Mogami T, Kon T, Yamamoto K. 2007. Kinetic mechanism of the fastest motor protein, Chara myosin. Journal of Biological Chemistry 282, 19534–19545.

Ito K, Yamaguchi Y, Yanase K, Ichikawa Y, Yamamoto K. 2009. Unique charge distribution in surface loops confers high velocity on the fast motor protein Chara myosin. *Proceedings of the National Academy of Sciences*, USA 106, 21585–21590.

Kakei T, Sumiyoshi H, Higashi-Fujime S. 2012. Characteristics of light chains of Chara myosin revealed by immunological investigation. Proceedings of the Japan Academy. Series B, Physical and Biological Sciences 88, 201–211.

Khan BR, Zolman BK. 2010. pex5 Mutants that differentially disrupt PTS1 and PTS2 peroxisomal matrix protein import in Arabidopsis. Plant Physiology 154, 1602–1615.

Kollmar, M., Mühlhausen, S. 2017. Myosin repertoire expansion coincides with eukaryotic diversification in the Mesoproterozoic era. BMC Evol Biol 17, 211. https://doi.org/10.1186/s12862-017-1056-2

Kumari P, Dahiya P, Livanos P, Zergiebel L, Kölling M, Poeschl Y, Stamm G, Hermann A, Abel S, Müller S, Bürstenbinder K. 2021. IQ67 DOMAIN proteins facilitate preprophase band formation and division-plane orientation. Nature Plants 7, 739–747.

Kurth EG, Peremyslov VV, Turner HL, Makarova KS, Iranzo J, Mekhedov SL., Koonin EV, Dolja VV. (2017). Myosin-driven transport network in plants. *Proceedings of the National Academy of Sciences*, USA 114, E1385–e1394.

Langelaan DN, Liburd J, Yang Y, Miller E, Chitayat S, Crawley SW, Côté GP, Smith SP. 2016. Structure of the Single-lobe myosin light chain c in complex with the light chain-binding domains of myosin-1c provides insights into divergent IQ motif recognition. Journal of Biological Chemistry 291, 19607–19617.

La Verde V, Dominici P, Astegno A. 2018. Towards understanding plant calcium signaling through calmodulin-like proteins: A biochemical and structural perspective. International Journal of Molecular Sciences 19, 1–18.

Lee E, Liu Z, Nguyen N, Nairn AC, Chang AN. 2022. Myosin light chain phosphatase catalytic subunit dephosphorylates cardiac myosin via mechanisms dependent and independent of the MYPT regulatory subunits. The Journal of Biological Chemistry 298, 102296.

Liu N, Lee L-Y, Hsu F-Y, Yu Y, Rao P, Gelvin SB. 2023. Myosin VIII and XI isoforms interact with Agrobacterium VirE2 protein and help direct transport from the plasma membrane to the perinuclear region during plant transformation. bioRxiv, 2023.03.06.531343.

Manceva S, Lin T, Pham H, Lewis JH, Goldman YE, Ostap EM. 2007. Calcium regulation of calmodulin binding to and dissociation from the myo1c regulatory domain. Biochemistry 46, 11718– 11726.

Ma YZ, Yen LF. 1989. Actin and Myosin in pea tendrils. Plant Physiology 89, 586–589.

McCormack E, Braam J. 2003. Calmodulins and related potential calcium sensors of Arabidopsis. The New Phytologist 159, 585–598.

McCurdy DW, Harmon AC. 1992. Phosphorylation of a putative myosin light chain inChara by calcium-dependent protein kinase. Protoplasma 171, 85–88.

Nebenführ A, Dixit R. 2018. Kinesins and myosins: molecular motors that coordinate cellular functions in plants. Annual Review of Plant Biology 69, 329–361.

Olatunji D, Kelley DR. 2020. A role for Arabidopsis myosins in sugar-induced hypocotyl elongation. MicroPublication Biology 2020.

Olatunji D, Clark NM, Kelley DR. 2022. The class VIII myosin ATM1 is required for root apical meristem function. bioRxiv.11.30.518567; doi: https://doi.org/10.1101/2022.11.30.518567

Paysan-Lafosse T, Blum M, Chuguransky S, et al. 2023. InterPro in 2022. Nucleic Acids Research 51, D418–D427.

Pazicky S, Dhamotharan K, Kaszuba K, Mertens HDT, Gilberger T, Svergun D, Kosinski J, Weininger U, Löw C. 2020. Structural role of essential light chains in the apicomplexan glideosome. Communications Biology 3, 568.

Peremyslov VV, Morgun EA, Kurth EG, Makarova KS, Koonin EV, Dolja VV. (2013). Identification of myosin XI receptors in *Arabidopsis* defines a distinct class of transport vesicles. The Plant Cell 25, 3022–3038.

Peremyslov VV, Prokhnevsky AI, Dolja VV. 2010. Class XI myosins are required for development, cell expansion, and F-Actin organization in Arabidopsis. The Plant Cell 22, 1883–1897.

Pitzalis N, Heinlein M. 2017. The roles of membranes and associated cytoskeleton in plant virus replication and cell-to-cell movement. Journal of Experimental Botany 69, 117–132.

Rula S, Suwa T, Kijima ST, Haraguchi T, Wakatsuki S, Sato N, Duan Z, Tominaga M, Uyeda TQP, Ito, K. (2018). Measurement of enzymatic and motile activities of Arabidopsis myosins by using Arabidopsis actins. Biochemical and Biophysical Research Communications 495, 2145–2151.

Sambrook J, Russell DW. 2001. *Molecular Cloning: A Laboratory Manual*.

Sattarzadeh A, Franzen R, Schmelzer E. (2008). The Arabidopsis class VIII myosin ATM2 is involved in endocytosis. Cell Motility and the Cytoskeleton 65, 457–468.

Shen M, Zhang N, Zheng S, Zhang W-B, Zhang H-M, Lu Z, Su QP, Sun Y, Ye K, Li X-D. 2016. Calmodulin in complex with the first IQ motif of myosin-5a functions as an intact calcium sensor. Proceedings of the National Academy of Sciences of the United States of America 113, E5812–E5820.

Sievers F, Higgins DG. 2014. Clustal omega. Current Protocols in Bioinformatics 48, 3.13.1-3.13.16.

Slaughter BD, Unruh JR, Allen MW, Bieber Urbauer RJ, Johnson CK. 2005. Conformational substates of calmodulin revealed by single-pair fluorescence resonance energy transfer: influence of solution conditions and oxidative modification. Biochemistry 44, 3694–3707.

Talts K, Ilau B, Ojangu EL, Tanner K, Peremyslov VV, Dolja VV, Truve E, Paves H. (2016). Arabidopsis Myosins XI1, XI2, and XIK are crucial for gravity-induced bending of inflorescence stems. Frontiers in Plant Science 7, 1932.

Teresinski HJ, Hau B, Symonds K, Kilburn R, Munro KA, Doner NM, Mullen R, Li VH, Snedden WA. 2023. Arabidopsis calmodulin-like proteins CML13 and CML14 interact with proteins that have IQ domains. bioRxiv, 2023.03.09.531943.

Tominaga M, Nakano A. 2012. Plant-Specific Myosin XI, a Molecular Perspective. Frontiers in Plant Science 3.

Vahey M, Titus M, Trautwein R, Scordilis S. 1982. Tomato actin and myosin: Contractile proteins from a higher land plant. Cell Motility 2, 131–147.

Vallone R, La Verde V, D’Onofrio M, Giorgetti A, Dominici P, Astegno A. 2016. Metal binding affinity and structural properties of calmodulin-like protein 14 from *Arabidopsis thaliana*. Protein Science 25, 1461–1471.

Van Leene J, Eeckhout D, Gadeyne A, Matthijs C, Han C, De Winne N, Persiau G, Van De Slijke E, Persyn F, Mertens T, Smagghe W, Crepin N, Broucke E, Van Damme D, Pleskot R, Rolland F, De Jaeger G. 2022. Mapping of the plant SnRK1 kinase signaling network reveals a key regulatory role for the class II T6P synthase-like proteins. Nature Plants 8, 1245–1261.

Vaughn JL, Goodwin RH, Tompkins GJ, McCawley P. 1977. The establishment of two cell lines from the insect *Spodoptera frugiperda* (lepidoptera; noctuidae). In Vitro 13, 213–217.

Vetter SW, Leclerc E. 2003. Novel aspects of calmodulin target recognition and activation. European journal of biochemistry. FEBS Letters 270, 404–414.

Wang Y, Wang B, Gilroy S, Wassim Chehab E, Braam J. 2011. CML24 is Involved in Root Mechanoresponses and Cortical Microtubule Orientation in Arabidopsis. Journal of Plant Growth Regulation 30, 467–479.

Wu S-Z, Bezanilla M. 2014. Myosin VIII associates with microtubule ends and together with actin plays a role in guiding plant cell division. eLife 3.

Wu S-Z, Ritchie JA, Pan A-H, Quatrano RS, Bezanilla M. 2011. Myosin VIII Regulates Protonemal Patterning and Developmental Timing in the Moss Physcomitrella patens. Molecular Plant 4, 909–921.

Yokota E, Muto S, Shimmen T. (1999). Inhibitory regulation of higher-plant myosin by Ca^2+^ ions. Plant Physiology 119, 231–240.

Yokota E, Shimmen T. 1994. Isolation and characterization of plant myosin from pollen tubes of lily. Protoplasma 177, 153–162.

Zhu X, Dunand C, Snedden W, Galaud J-P. 2015. CaM and CML emergence in the green lineage. Trends in Plant Science 20, 483–489.

